# Evaluating material-driven regeneration in a tissue engineered human *in vitro* bone defect model

**DOI:** 10.1101/2022.08.05.502914

**Authors:** Bregje W.M. de Wildt, Esther E.A. Cramer, Leanne S. de Silva, Keita Ito, Debby Gawlitta, Sandra Hofmann

## Abstract

Advanced *in vitro* human bone defect models can contribute to the evaluation of materials for *in situ* bone regeneration, addressing both translational and ethical concerns regarding animal models. In this study, we attempted to develop such a model to study material-driven regeneration, using a tissue engineering approach. By co-culturing human umbilical vein endothelial cells (HUVECs) with human bone marrow-derived mesenchymal stromal cells (hBMSCs) on silk fibroin scaffolds with *in vitro* critically sized defects, the growth of vascular-like networks and three-dimensional bone-like tissue was facilitated. After a model build-up phase of 28 days, materials were artificially implanted and HUVEC and hBMSC migration, cell-material interactions, and osteoinduction were evaluated 14 days after implantation. The materials physiologically relevant for bone regeneration included a platelet gel as blood clot mimic, cartilage spheres as soft callus mimics, and a fibrin gel as control. Although the *in vitro* model was limited in the evaluation of immune responses, hallmarks of physiological bone regeneration were observed *in vitro*. These included the endothelial cell chemotaxis induced by the blood clot mimic and the mineralization of the soft callus mimic. Therefore, the present *in vitro* model could contribute to an improved pre-clinical evaluation of biomaterials while reducing the need for animal experiments.

**Graphical abstract:** 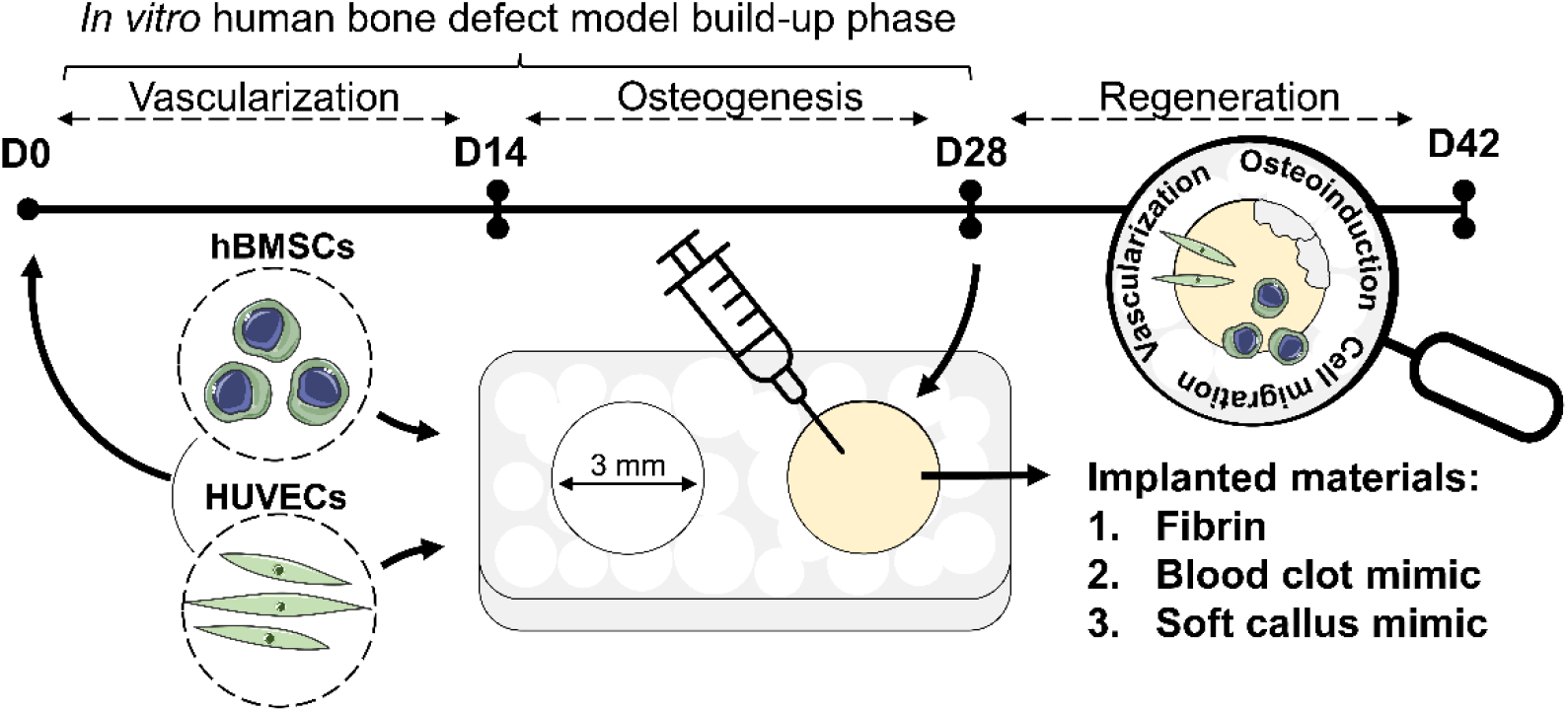

## 1. Introduction

Bone is a highly vascularized and dynamic tissue with the capacity to regenerate without scar formation. Nevertheless, in 2% – 5% of the defects, the failure of bone to bridge the gap results in a non-union [1,2]. Bone tissue engineering has been focusing on the development of implantable grafts to tackle such defects. While initially grafts were grown prior to implantation *in vitro* by making use of biomaterials, progenitor cells, and stimuli, current grafts are more often developed to be intelligent biomaterials for *in situ* regeneration, making use of the bone’s innate capacity to regenerate upon implantation [3,4]. After successful *in vitro* assessments, these grafts are routinely studied in animal models [5–7]. Despite animal studies being a crucial step in elucidating material-host interactions, the translation from *in vitro* to *in vivo* experiments has been poor. The current pace at which materials are being developed causes a significant burden on animal experiments [8]. Moreover, with a success rate of less than 10%, the subsequent clinical translation of *in vivo* animal models is also poor [9,10], which is likely caused by their insufficient representation of the human physiology [11]. Thus, both the translation from *in vitro* assessments to *in vivo* models and the translation from *in vivo* animal models to the human clinic need to be improved. To address these translational issues and improve preclinical graft testing, advanced human *in vitro* bone defect models should be developed and integrated into the preclinical graft testing routine [12–14]. For the creation of such *in vitro* models, traditional bone tissue engineering strategies can be applied [15].

While bone tissue engineering strategies have already been successfully applied for the creation of *in vitro* models for bone marrow [16], bone metastasis [17], woven bone [18] and bone remodeling [19], the development of human *in vitro* bone defect models for biomaterial testing is rarely explored [14,15]. A tissue engineered defect model has previously been proposed, where authors created defects in silk fibroin (SF) scaffolds, seeded scaffolds with human bone marrow-derived mesenchymal stromal cells (hBMSCs), and studied tissue growth and mineralization in the defect area upon osteogenic differentiation [20]. In another study, the *in vitro* evaluation of biomaterial-induced bone regeneration was analyzed using a fibrin matrix that was sandwiched between two human trabecular bone discs loaded with rabbit periosteal cells [21]. Constructs were subsequently mechanically stimulated and histologically evaluated after 14 days of culture [21]. Although they observed osteogenic differentiation of the periosteal cells under influence of mechanical loading, cell migration into the defect site was not detected.

Successful material-driven bone regeneration relies on a cascade of biological events, including: (i) inflammation and immunomodulation, (ii) progenitor cell migration and differentiation, (iii) vascularization, (iv) osteoinduction, (v) implant remodeling [4]. As such, models aiming at recapitulating material-driven bone regeneration *in vitro* should be able to capture such events. Especially the materials’ ability to stimulate vascularization is of interest as this is crucial for successful fracture healing [22], and it is therefore the current major challenge in regenerative treatments of large bone defects [23].

In this study, we attempted to develop an *in vitro* human bone defect model to study material-driven regeneration, using a tissue engineering approach (Figure 1A). To enable the evaluation of a material’s potential to stimulate vascularization, cell migration and osteoinduction, hBMSCs were co-cultured with human umbilical vein endothelial cells (HUVECs) to generate the bone compartment containing the defects (Figure 1A). In a 3-dimensional (3D) microenvironment, HUVECs are capable of forming vascular networks in co-culture with hBMSCs; where hBMSCs function as supporting cells by surrounding vessel-like structures and providing angiogenic factors [24,25]. In turn, endothelial cells support osteogenic differentiation by the secretion of factors like bone morphogenetic proteins [26]. As a result, hBMSCs can additionally undergo osteogenic differentiation in the 3D space between the capillary-like network [27,28]. To facilitate 3D growth of these cells, a SF scaffold was used with two *in vitro* critically sized defects of 3 mm in diameter, as based on previous experiments [20] (Figure 1A). First, the bone defect model was created by 14 days of culture to stimulate vascularization, followed by 14 days of culture to stimulate osteogenic differentiation and bone-like matrix formation (Figure 1B). At day 28, defects were filled with physiologically relevant graft materials, aiming at mimicking physiological bone regeneration for the validation of our model (Figure 1C). Implanted materials included a human platelet lysate gel as blood clot mimic, devitalized cartilage spheres as soft callus mimics [29], and a fibrin gel as control. By tracking the defect model with non-destructive confocal microscopy, the materials’ potential to stimulate cell migration, vascularization and osteoinduction were evaluated.

**Figure 1.**
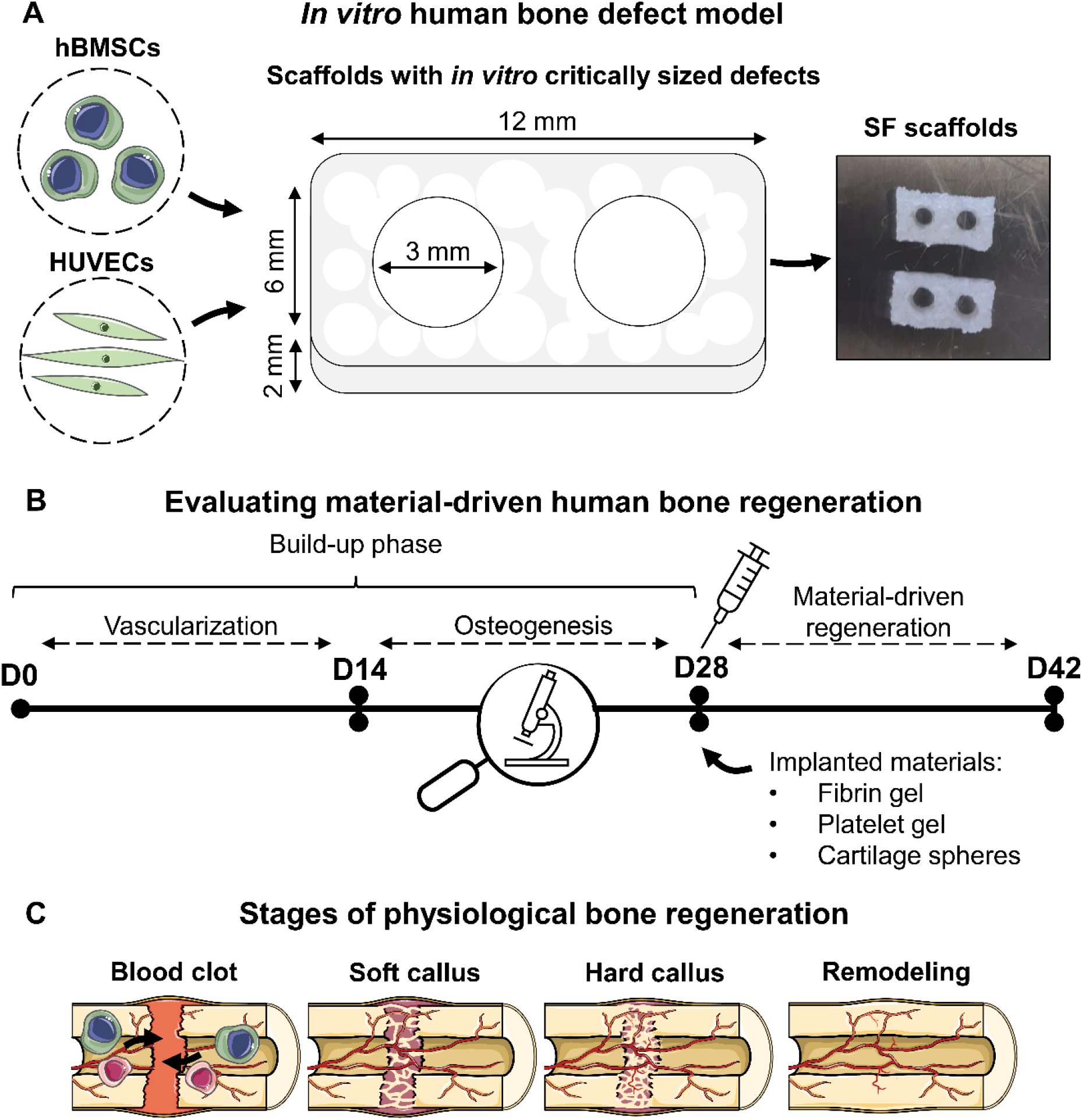
Experimental set-up of the presented study. (**A**) A co-culture of hBMSCs and HUVECs was performed on SF scaffolds with two critically sized defects. (**B**) To create the *in vitro* bone defect model, cells were first stimulated to form vascular-like networks for 14 days. hBMSCs were subsequently stimulated to undergo osteogenic differentiation and produce bone-like matrix for 14 days. After 28 days, materials were implanted, and regeneration was studied after an additional 14 days of culture (day 42). Fibrin gel, platelet gel as blood clot mimic, and cartilage spheres as soft callus mimics were implanted, aiming at capturing physiological stages of bone regeneration (**C**). The figure was modified from Servier Medical Art, licensed under a Creative Common Attribution 3.0 Generic License (http://smart.servier.com/, accessed on 20 May 2022). Abbreviations: human bone marrow-derived mesenchymal stromal cells (hBMSCs), human umbilical vein endothelial cells (HUVECs), silk fibroin (SF), day (D).

## 2. Materials and Methods

### 2.1 Scaffold fabrication

Bombyx mori L. silkworm cocoons were degummed by boiling them in 0.2 M Na2CO3 (S-7795, Sigma-Aldrich, Zwijndrecht, The Netherlands) for 1 h. After drying, silk was dissolved in 9 M LiBr (199870025, Acros, Thermo Fisher Scientific, Breda, The Netherlands), filtered, and dialyzed against ultra-pure water (UPW) for 36 h using SnakeSkin Dialysis Tubing (molecular weight cut-off: 3.5 K, 11532541, Thermo Fisher Scientific). The dialyzed SF solution was frozen at -80° C and lyophilized for 7 days. Lyophilized SF was dissolved in hexafluoro-2-propanol (003409, Fluorochem, Hadfield, UK) at a concentration of 17% (w/v) and casted in scaffold molds containing NaCl granules with a size of 425-500 µm as template for the pores. Molds were covered to improve the SF blending with the granules. After 3 h, covers were removed from molds, and hexafluoro-2-propanol was allowed to evaporate for 7 days whereafter *β-* sheets were induced by submerging SF-salt blocks in 90% MeOH for 30 min. SF-salt blocks were cut into discs of 2 mm height with a Accutom-5 (04946133, Struer, Cleveland, OH, USA). NaCl was dissolved for 48 h from the scaffolds in UPW, resulting in porous sponges. These sponges were cut into scaffolds with a length of 12 mm and a width of 6 mm and provided with two central defects with a 3 mm diameter biopsy punch. The dimensions of the defects were based on previous research in which a 3 mm channel remained open over a period of 42 days [20]. Scaffolds were sterilized by autoclaving in phosphate buffered saline (PBS) at 121° C for 20 min.

### 2.2 Cell culture experiments

#### 2.2.1 hBMSC isolation, expansion and seeding

Mesenchymal stromal cells were isolated from human bone marrow (1M-125, Lonza, Walkersville, MD, USA, collected under their institutional guidelines and with informed consent) and characterized for surface markers and multilineage differentiation, as previously described [30]. hBMSCs were frozen at passage 4 with 5*10^6^ cells/ml in freezing medium containing fetal bovine serum (FBS, BCBV7611, Sigma-Aldrich) with 10% DMSO and stored in liquid nitrogen until further use. Before experiments, hBMSCs were thawed, collected in high glucose DMEM (hg-DMEM, 41966, Thermo Fisher Scientific), seeded at a density of 2.5*10^3^ cells/cm^2^ and expanded in expansion medium containing hg-DMEM, 10% FBS (BCBV7611, Sigma-Aldrich), 1% Antibiotic Antimycotic (anti-anti, 15240, Thermo Fisher Scientific), 1% Non-Essential Amino Acids (11140, Thermo Fisher Scientific), and 1 ng/ml basic fibroblast growth factor (bFGF, 100-18B, PeproTech, London, UK) at 37 °C and 5% CO_2_. After 10 days at around 80% confluence, cells were detached using 0.25% trypsin-EDTA (25200, Thermo Fisher Scientific) and seeded onto scaffolds at passage 5. Cells were seeded at a density of 1*10^6^ cells per scaffold and seeding was performed dynamically [31] for 6 hours in 50 ml tubes on an orbital shaker at 150 RPM in osteogenic control medium (low glucose-DMEM (lg-DMEM, 22320, Thermo Fisher Scientific), 10% FBS (SFBS, Bovogen, East Keilor, Australia) and 1% anti-anti). After seeding, scaffolds were transferred to 24-wells plates and kept overnight in endothelial cell growth medium-2 (EGM-2, CC-3162, Lonza). HUVECs were seeded the next day.

#### 2.2.2 HUVEC expansion and seeding

Pooled primary green fluorescent protein (GFP) expressing HUVECs (GFP, Olaf Pharmaceuticals, Worcester, MA, USA) were expanded in EGM-2 with 3% extra FBS (FBS, BCBV7611, Sigma-Aldrich) to passage 8. After 5 days, HUVECs were detached using 0.25% trypsin-EDTA. Just before seeding, EGM-2 was removed from hBMSC-containing scaffolds. HUVECs were resuspended at a concentration of 1*10^6^/50 µl and cells were seeded with a 50 µl drop onto the hBMSC-containing scaffolds. Scaffolds were incubated at 37 °C for 20 min to allow for cell attachment, whereafter 2 ml EGM-2 was added to the wells which is referred to as day 0 of the experiment.

#### 2.2.3 hBMSC-HUVEC co-culture

Constructs (*N* = 16) were incubated for 42 days at 37 °C and 5% CO_2_ and initially provided with EGM-2 to allow for the development of vascular-like networks. After 14 days, when vascular-like structures were formed, medium was switched to osteogenic differentiation medium (osteogenic control medium with osteogenic supplements 10 mM *β*-glycerophosphate (G9422, Sigma-Aldrich), 50 µg/ml ascorbic acid-2-phosphate (A8960, Sigma Aldrich), and 100 nM Dexamethasone (D4902, Sigma-Aldrich)) for the remaining culture period. During the whole experiment, medium was changed 3 times a week. On day 28, some scaffolds (*N* = 4) were sacrificed to assess osteogenic differentiation and mineralization. Other scaffolds were kept in culture for another 14 days.

#### 2.2.4 Preparation of devitalized cartilage spheres

Cartilaginous spheres as soft callus mimics were grown from hBMSCs as previously described [29]. Briefly, 20*10^6^ hBMSCs were encapsulated in collagen type I gel droplets (4 mg/ml) (354249, Corning, New York, USA), according to the manufacturer’s instructions. After gelation, samples were cultured in chondrogenic differentiation medium (hg-DMEM (31966, Thermo Fisher Scientific), 1% insulin-transferrin-selenium + premix (354352, Corning), 100 nM dexamethasone (D8893, Sigma-Aldrich), 50 µg/ml ascorbic acid-2-phosphate, 100 units/ml of penicillin and 100 µg/ml streptomycin (15140, Thermo Fisher Scientific), and 10 ng/ml transforming growth factor-*β*1 (Peprotech). Spheres were cultured for 28 days. For the first 4 days, medium was refreshed daily and afterwards 3 times per week. After 28 days, spheres were devitalized by a mild procedure and stored frozen until use.

#### 2.2.5 Material implantation

After 28 days of culture, three different materials were artificially implanted: (i) a human fibrin gel, (ii) a human platelet lysate gel as blood clot mimic, and (iii) two devitalized cartilage spheres as soft callus mimics. Just before implantation, medium was removed of all scaffolds. For the fibrin gel, fibrinogen (341576-M, Sigma-Aldrich) was mixed with thrombin (T6884, Sigma-Aldrich) to a final concentration of 2.5 mg/ml fibrinogen and 0.2 U/ml thrombin. In each defect (*N* = 4 scaffolds and *N* = 8 defects), 50 µl fibrin was pipetted, whereafter samples were incubated at 37 °C for 25 min to allow for polymerization of the gel. For the platelet gel, ELAREM™ matrix kit (MA30311) was used according to the manufacturer’s instructions. A 10% ELAREM™ matrix solution was prepared in PBS to which 50 µg/ml ELAREM™ accelerator was added. In each defect (*N* = 4 scaffolds and *N* = 8 defects), 50 µl platelet gel was pipetted, whereafter samples were placed in the incubator at 37 °C for 2 min to allow for gelation. For the cartilage spheres, spheres were soaked for 5 min in lg-DMEM to allow for rehydration of the spheres. Meanwhile, fibrin gels were prepared and 25 µl was implanted as described above. Before polymerization of the gel, two spheres per defect (*N* = 4 scaffolds and *N* = 8 defects) were dried with a sterile gauze and implanted. After implantation, a small droplet of fibrin gel was added on top of the spheres to fill the defect. Samples were incubated at 37 °C for 25 min to allow for polymerization of the gel. After proper gelation in all conditions, osteogenic differentiation medium was added, and defect regeneration was followed after an additional 14 days of culture.

### 2.3 *In vitro* model analyses

#### 2.3.1 Live confocal microscopy

During the culture period, vascularization (day 4, 7, 11 and 14) (*N* = 16 scaffolds), bone-like matrix production (day 28) (*N* = 4 scaffolds), and cell migration (day 35 and 42) (*N* = 4 defects per condition) were visualized with microscopy. HUVECs were visualized by their GFP-label. Collagen, hydroxyapatite and hBMSCs were visualized with viable dyes from day 28 on. On day 27 (end-point samples), 34 and 41, samples were washed in lg-DMEM and samples were stained overnight at 37 °C in osteogenic control medium (Section 2.2.1) with 0.2 nmol/ml OsteoSense™ 680 (dissolved in PBS; NEV10020EX, PerkinElmer, Waltham, MA, USA) to visualize hydroxyapatite, and 1 μmol/ml in-house made CNA35-mCherry [32] to visualize collagen. The next day, one droplet of NucBlue™ Hoechst 33342 (R37605, Thermo Fisher Scientific) was added per scaffold to visualize all cell nuclei and samples were incubated at 37 °C for 20 min. Samples were washed three times in lg-DMEM and provided with fresh osteogenic differentiation medium (Section 2.2.3). Data were acquired on a confocal laser scanning microscope equipped with a multi-photon laser and incubation unit (Leica TCS SP8X, 10x/0.40 HC PL Apo CS2 objective). During imaging, samples were kept at 37 °C and 5% CO_2_. To visualize cell nuclei, the multiphoton laser was used at *λ* = 740 nm to reduce cytotoxicity that can be induced by using violet light [33]. To study the influence of staining on cell death, unstained and stained samples were additionally compared for their lactate dehydrogenase (LDH) release in the supernatant (Section 2.3.5)

#### 2.3.2 Micro-computed tomography (*µ*CT)

After 42 days, scaffold halves (*N* = 4 per condition) were fixed in 3.7% neutral buffered formaldehyde overnight. Samples were scanned and analyzed with a *µ*CT100 imaging system. Scanning was performed at an isotropic nominal resolution of 17.2 µm, energy level of 45 kVp, intensity of 200 µA, integration time of 300 ms and with twofold frame averaging. To reduce part of the noise, a constrained Gaussian filter was applied with filter support 1 and filter width sigma 0.8 voxel. Filtered images were contoured for the scaffold and its 3 mm diameter defect and segmented to detect mineralization at a global threshold of 24% of the maximum grayscale value. To further reduce noise, unconnected objects smaller than 30 voxels were removed through component labeling. Mineralized volumes were computed for the total scaffold and the defect region using the scanner manufacturer’s image processing language (IPL) [34].

#### 2.3.3 Biochemical content analysis

To quantify the biochemical content, scaffolds were cut in halves (*N* = 4 per condition) and their defect content was collected by punching with a 3 mm biopsy punch. Scaffolds and defects were lyophilized, dry weights of scaffolds were collected and samples were digested overnight in papain digestion buffer (containing 100 mmol phosphate buffer, 5 mmol L-cystein, 5 mmol EDTA and 140 µg/ml papain (P4762, Sigma-Aldrich)) at 60 °C. DNA content of scaffolds and defects was quantified using the Qubit Quantification Platform (Invitrogen) with the high sensitivity assay, according to the manufacturer’s instructions. Hydroxyproline content as a measure for collagen was quantified in scaffolds using a chloramine-T assay [35] with trans-4-hydroxyproline (H54409, Sigma-Aldrich) as reference. Absorbance values were measured at 550 nm using a plate reader (Synergy™ HTX, Biotek) and standard curve absorbance values were used to determine hydroxyproline content in the samples. Glycosaminoglycan (GAG) content of scaffolds was measured using a dimethyl methylene blue (DMMB) assay [36] with shark cartilage chondroitin sulfate (C4284, Sigma-Aldrich) as a reference. Absorbance was read at 540 nm and 595 nm using a plate reader. Absorbance values were subtracted from each other (540-595) and converted to GAG content using standard curve absorbance values.

#### 2.3.4 (Immuno)histochemical analyses

Scaffolds halves (*N* = 4 per condition) were prepared for cryosections by soaking them for 15 minutes in each 5% (w/v) sucrose and 35% (w/v) sucrose in PBS. Samples were embedded in Tissue Tek® (Sakura), quickly frozen with liquid N_2_ and cryosections were prepared. Upon staining, sections were fixed in 3.7% neutral buffered formaldehyde and washed twice with PBS. To visualize collagen deposition, 5 μm thick cryosections (*N* = 4 scaffolds per condition) were stained with Picrosirius Red. Sections were soaked in Weigert’s Iron Hematoxylin (HT1079, Sigma-Aldrich) solution for 10 minutes, washed in running tap water for 10 minutes, and stained in 1% w/v Sirius Red (36,554-8, Sigma-Aldrich) in picric acid solution (36011, Sigma-Aldrich) for one hour. Subsequently, sections were washed in two changes of 0.5% acetic acid and dehydrated in one change of 70% and 96% EtOH, three changes of 100% EtOH, and two changes of xylene. Sections were mounted with Entellan (107961 Sigma-Aldrich). To capture the entire sample, tile scans were made with a bright field microscope (Zeiss Axio Observer Z1, 10x/0.45 Plan-Apochromat objective). Tile scans were stitched with Zen Blue software (version 3.1, Zeiss).

To study osteogenic differentiation, 5 μm thick cryosections (*N* = 2 scaffolds per condition) were stained with 1 µg/ml DAPI and antibodies for runt-related transcription factor-2 (RUNX2) and osteopontin. Collagen deposition was characterized by staining 5 μm thick cryosections (*N* = 2 scaffolds per condition) with 1 µg/ml DAPI and a collagen type I antibody. To study the cell-material interactions and the supporting cell functionality of the hBMSCs, 30 μm thick cryosections (*N* = 4 scaffolds of the cartilage spheres group) were stained with 1 µg/ml DAPI, 50 pmol Atto 488-conjugated Phalloidin (49409, Sigma-Aldrich) and antibodies for CD31 and *α*-smooth muscle actin. Briefly, sections were permeabilized in 0.5% Triton X-100 in PBS (5 min for 5 μm sections and 10 min for 30 μm sections) and blocked in 10% normal goat serum in PBS for 30 min. Primary antibodies were incubated overnight at 4 °C in 1% normal goat serum in PBS, secondary antibodies were incubated with DAPI and Phalloidin (if applicable) for 1 h at room temperature. Antibodies are listed in Table S1. Images of the osteogenic differentiation and collagen type I staining were acquired with an epi-fluorescence microscope (Zeiss Axio Observer 7, 20x/0.4 LD Plan-Neofluor objective), and tile scans were stitched with Zen Blue software (version 3.3, Zeiss, Breda, The Netherlands). Z-stacks of 30 μm thick sections to visualize cell-material interactions were acquired with a laser scanning microscope (Leica TCS SP8X, 40x/0.95 HC PL Apo objective). Z-stacks were converted to maximum intensity projections using FiJi [37].

#### 2.3.5 LDH activity

To evaluate potential cytotoxic effects of implanted materials, LDH activity was measured in the cell supernatant on day 42 (*N* = 4 per condition). A 100 µl supernatant sample or NADH (10107735001, Sigma-Aldrich) standard was incubated with 100 µl LDH reaction mixture (11644793001, Sigma-Aldrich) in 96-wells assay plates. Absorbance was measured directly after the reaction mixture was added and after 30 min at 492 nm. LDH activity was calculated between the initial absorbance values and the absorbance values after 30 min reaction, using standard curve absorbance values.

#### 2.3.6 Multiplex immunoassays

To evaluate the protein content in the cell supernatant on day 42 (*N* = 4 per condition), a total of 21 proteins as markers for cell migration, vascularization, remodeling and bone formation were quantified using multiplex immunoassays at the Multiplex Core Facility (MCF) of the Laboratory for Translational Immunology of the University Medical Center Utrecht, the Netherlands. Immunoassays were developed and validated by the MCF and based on Luminex xMap technology (Luminex, Austin, TX, USA) [35]. Briefly, samples were incubated with MagPlex microspheres (Luminex) for 1 h at room temperature with continuous shaking, followed by 1 h incubation with biotinylated antibodies and 10 min incubation with phycoerythrin-conjugated streptavidin in high performance ELISA buffer (HPE, Sanquin, Hamburg, Germany). Data acquisition was performed with FLEXMAP 3D equipment in combination with xPONENT software (version 4.3, Luminex), and analyzed by 5-parametric curve fitting using Bio-Plex Manager software. Protein concentrations were normalized by converting them into z-scores (*i.e*., the number of standard deviations from the overall sample average) and presented using Heatmapper [36].

### 2.4 Statistical analyses

Statistical analyses were performed, and graphs were prepared in GraphPad Prism (version 9.3.0, GraphPad, La Jolla, CA, USA) and R (version 4.1.2) [37]. Data were tested for normality in distributions and equal variances using Shapiro-Wilk tests and Levene’s tests, respectively. When these assumptions were met, mean ± standard deviation are presented, and to test for differences, a one-way ANOVA was performed followed by Holm-Šídák’s method with adjusted *p*-values for multiple comparisons. Other data are presented as median ± interquartile range and were tested for differences with a non-parametric Kruskal-Wallis test followed by Dunn’s tests with adjusted *p*-values for multiple comparisons. A *p*-value of <0.05 was considered statistically significant.

## 3. Results

### 3.1 Creation of a tissue engineered human *in vitro* bone defect model

By microscopically evaluating GFP-labeled HUVECs in co-culture with hBMSCs during the initial 14 days of culture, the development of vascular-like structures after 14 days under the influence of EGM-2 medium was confirmed (Figure 2). On day 4 and 7, HUVECs appeared more as single cells that attached to the scaffold wall concavities where they started to form circular-like networks. From day 11 onwards, clear vascular-like networks with a branched morphology were found which started forming tubular structures (Figure 2), white arrows). HUVECs and vascular-like structures appeared throughout the whole scaffolds, which can be appreciated from the defect overview images (Figure 2).

**Figure 2.**
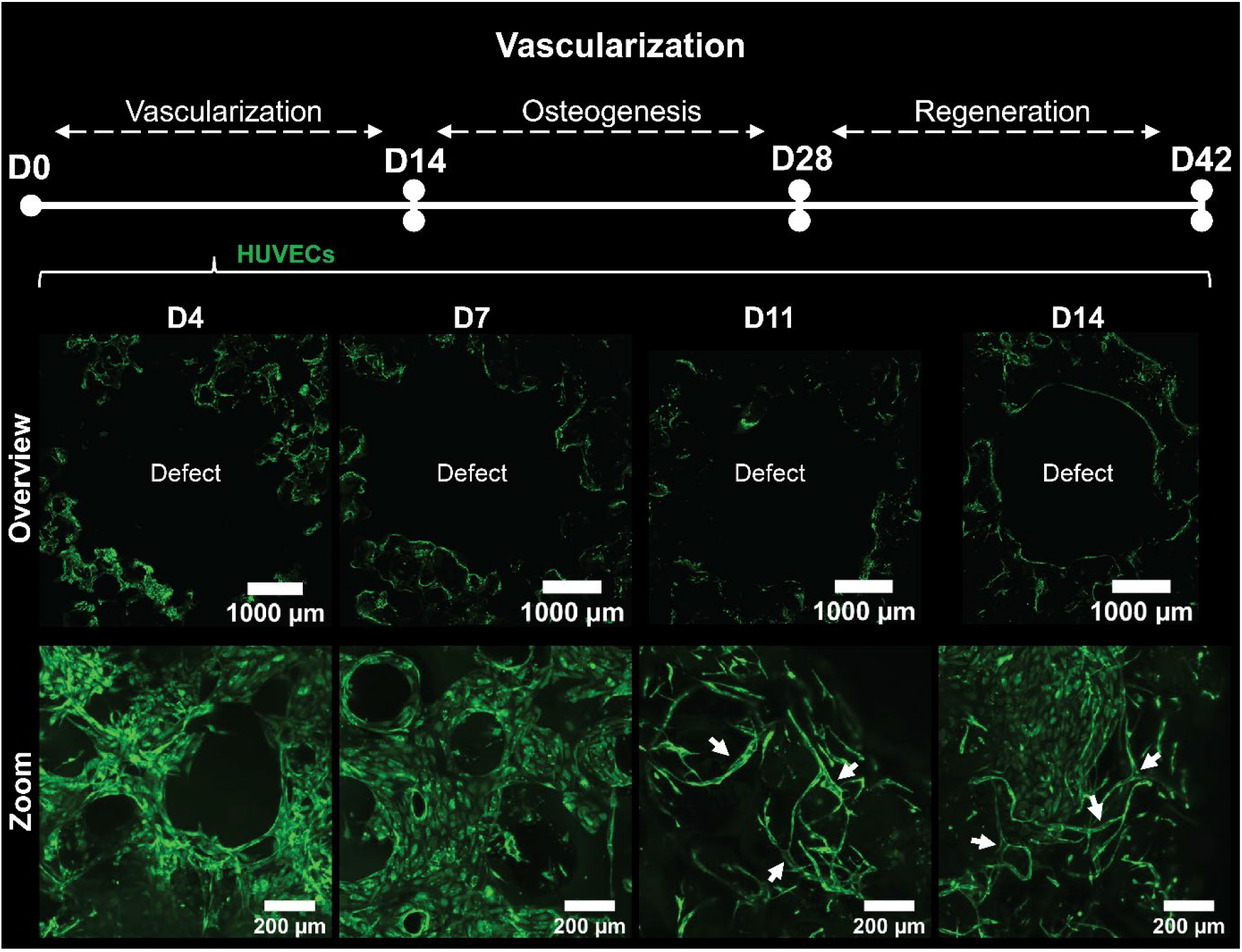
Microscopic evaluation of GFP-expressing HUVECs during the initial 14 days of culture to evaluate vascular-like network development. In the top panel, defect overview images are presented. At the bottom panel, close-up z-stack maximum intensity projection images are presented. White arrows indicate tubular vascular networks. Abbreviations: green fluorescent protein (GFP), human umbilical vein endothelial cells (HUVECs), day (D).

After 14 days, medium was switched to osteogenic differentiation medium to induce bone-like matrix formation and osteogenic differentiation of hBMSCs. Indeed, after an additional 14 days (*i.e*., day 28 of culture) hBMSCs had produced a bone-like extracellular matrix as observed from the collagen and hydroxyapatite stainings (Figure 3A), which are the two main components of the bone extracellular matrix (106). Staining of samples did not induce additional cell death (Figure S1). The defect of the scaffolds remained unfilled, confirming its critical size for *in vitro* experiments. Both single HUVECs and HUVECs organized in vascular-like networks were observed after 28 days (Figure 3A and Video S1). The presence of single HUVECs indicates that some vascular-like networks might have regressed after the switch from endothelial growth medium to osteogenic differentiation medium. Mineralization of the bone defect model was also confirmed by *µ*CT imaging, showing mineralized matrix throughout the whole scaffold (Figure 3B). By immunohistochemical analyses, collagen type I production was confirmed (Figure 3C). Osteogenic differentiation of hBMSCs was observed by the presence of the nuclear transcription factor RUNX2 and the non-collagenous protein osteopontin in the proximity of RUNX2 positive cells, as typical markers for osteogenesis (Figure 3D) [38]. As a next step, materials were implanted into this bone-like defect model possessing vascular-like networks.

**Figure 3.**
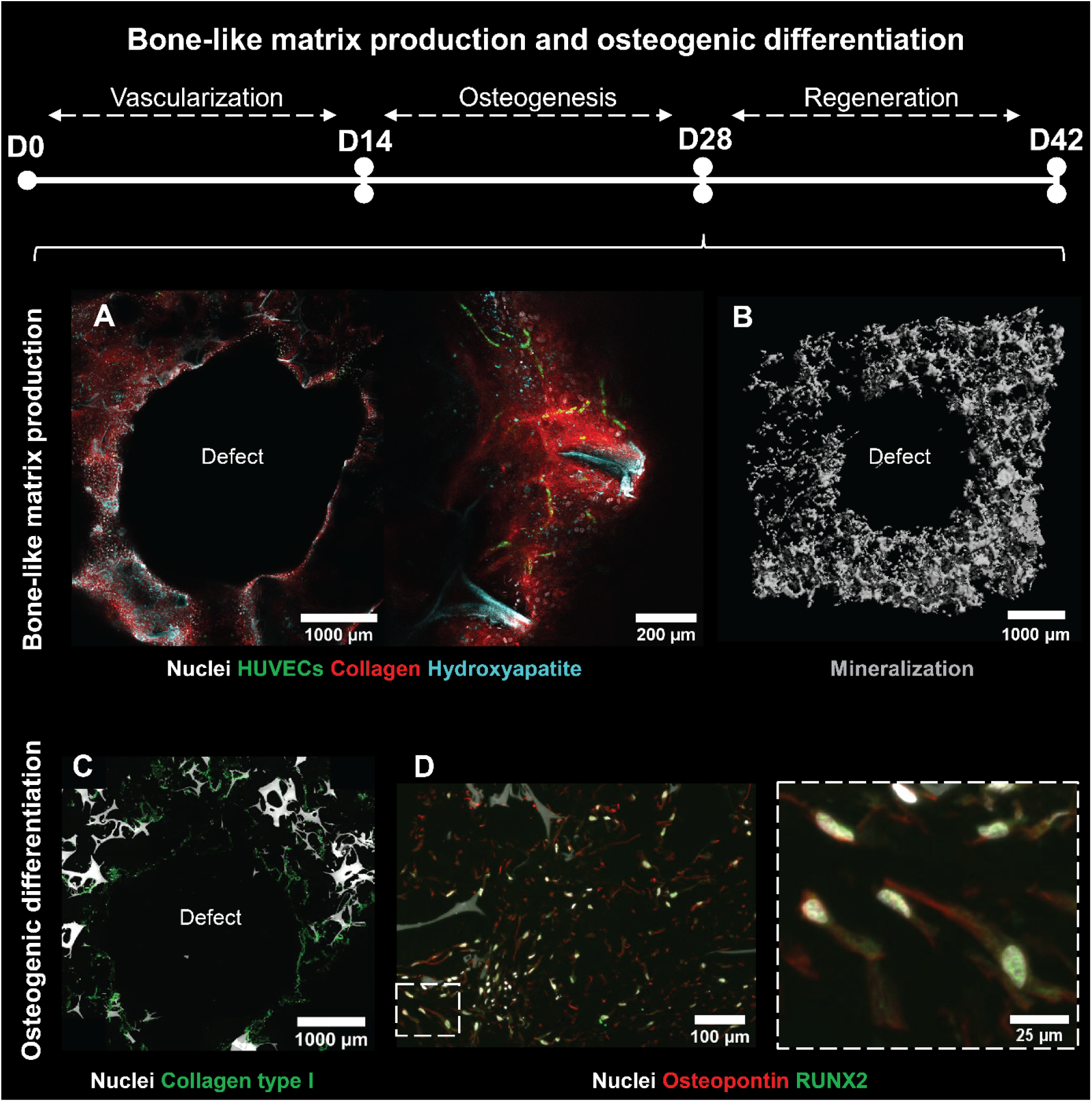
Evaluation of the osteogenesis phase after 28 days of culture. (**A**) Defect overview image (left) and close-up image (right) of viable construct containing GFP-expressing HUVECs (green) and stained for nuclei (gray), collagen (red), and hydroxyapatite (cyan). (**B**) *µ*CT reconstruction of scaffolds indicating mineralization through the whole defect model. (**C**) Collagen type I (green), (**D**) osteopontin (red), and RUNX2 (green) immunohistochemical analyses indicated osteogenic differentiation of the hBMSCs in the model. Abbreviations: green fluorescent protein (GFP), human umbilical vein endothelial cells (HUVECs), day (D), micro-computed tomography (*µ*CT), runt-related transcription factor-2 (RUNX2), human bone marrow-derived mesenchymal stromal cells (hBMSCs). (The reader is referred to the web version of this article for colored images)

### 3.2 Cell migration into the defect area upon material implantation

Upon material implantation, cell migration was microscopically evaluated on day 35 and 42. On day 35, defects implanted with fibrin gel or platelet gel showed little to no cell migration (Figure 4A+B). Only defects implanted with cartilage spheres showed clear migration of both HUVECs and (likely osteogenically differentiated) hBMSCs (Figure 4C), green cells and white nuclei, respectively) into the fibrin around the spheroids, already at day 35. Interestingly, HUVECs appeared to have migrated to the defect wall of defects implanted with platelet gel (Figure 4B). In contrast, this was not observed in defects implanted with fibrin (Figure 4A), indicating that the HUVECs require a chemical stimulus, likely present in the platelet gel, to migrate. These observations were confirmed after 42 days of culture. hBMSCs migrated to the implanted material in all conditions, but in defects implanted with platelet gel and cartilage spheres, also HUVECs migrated which was not observed in defects implanted with fibrin gel (Figure 4D-F, Video S2-S4 and Figure S2 for overview images). The migration of these cells appeared only in distinct areas (Figure S2), which could indicate the degradation or contraction of the implanted gels as observed before [39]. Defects implanted with cartilage spheres remained stable over time. Cell migration of both hBMSCs and HUVECs was observed around the spheres and HUVECs even attached to the cartilage spheres (Figure 4F and Video S4).

**Figure 4.**
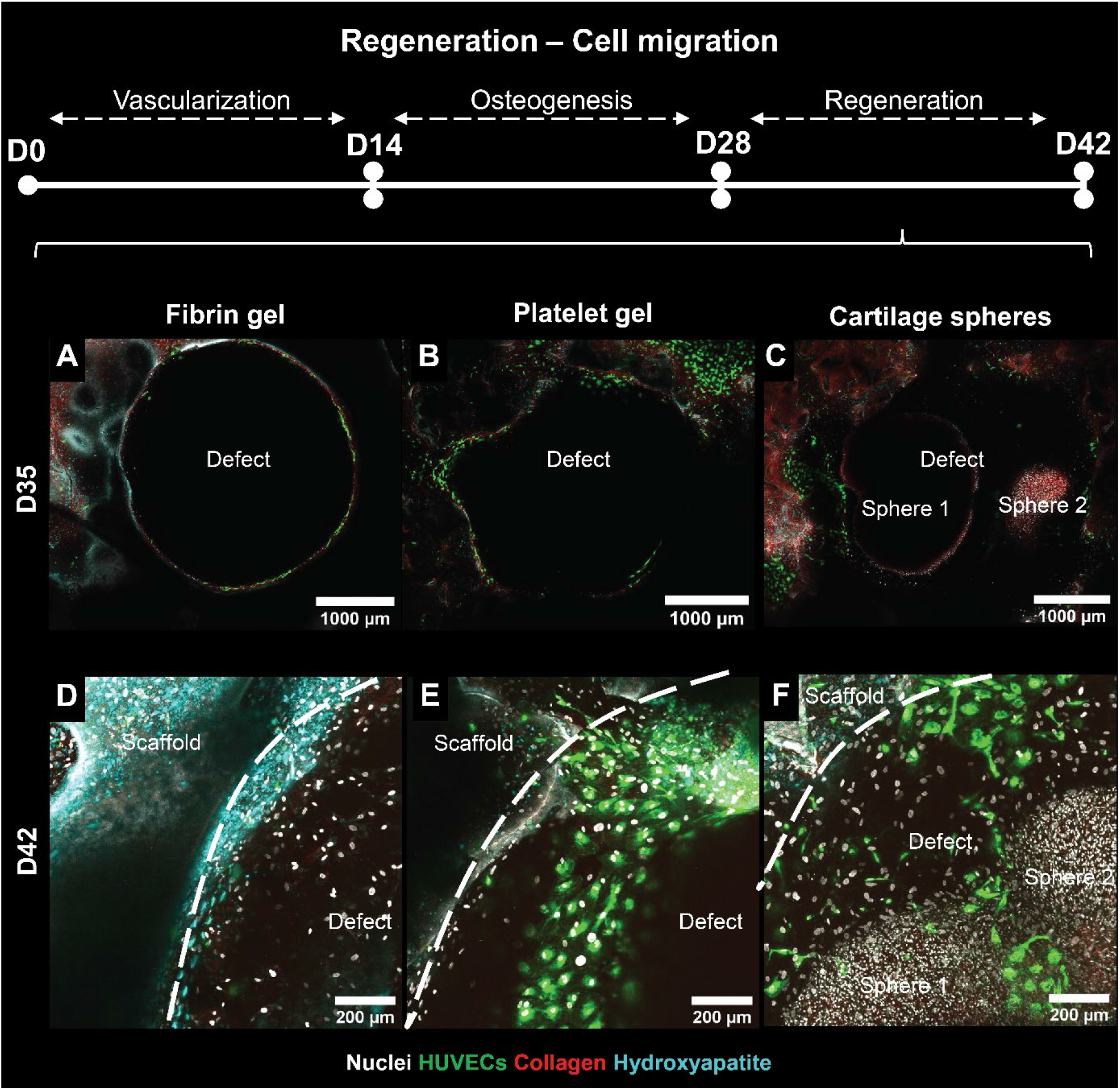
Evaluation of the regeneration phase of viable constructs after 35 days and 42 days of culture. Top panel presents defect overview images at day 35, bottom panel presents close-up z-stacks maximum intensity projection images at day 42 of culture of construct containing GFP-expressing HUVECs (green) and stained for nuclei (gray), collagen (red), and hydroxyapatite (cyan), implanted with (**A+D**) fibrin gel, (**B+E**) platelet gel, or (**C+F**) two cartilage spheres. Abbreviations: green fluorescent protein (GFP), human umbilical vein endothelial cells (HUVECs), day (D). (The reader is referred to the web version of this article for colored images)

### 3.3 Cell-material interactions

To evaluate the interactions of the cells with the implanted materials, cell culture medium supernatants of day 42 were analyzed for their protein content (after 2 days incubation with the samples). Protein concentrations were converted to Z-scores (*i.e*. the normalized deviation from the average of all experimental groups) and color-coded for presentation (Figure 5A). Of importance, these proteins can either be secreted by the cells or be a product of material degradation. Interestingly, all 21 studied proteins were detected in the culture medium supernatants at concentrations above the concentration measured in control medium (*i.e*., fresh osteogenic differentiation medium that has not been in contact with the cells) (Table S2).

**Figure 5.**
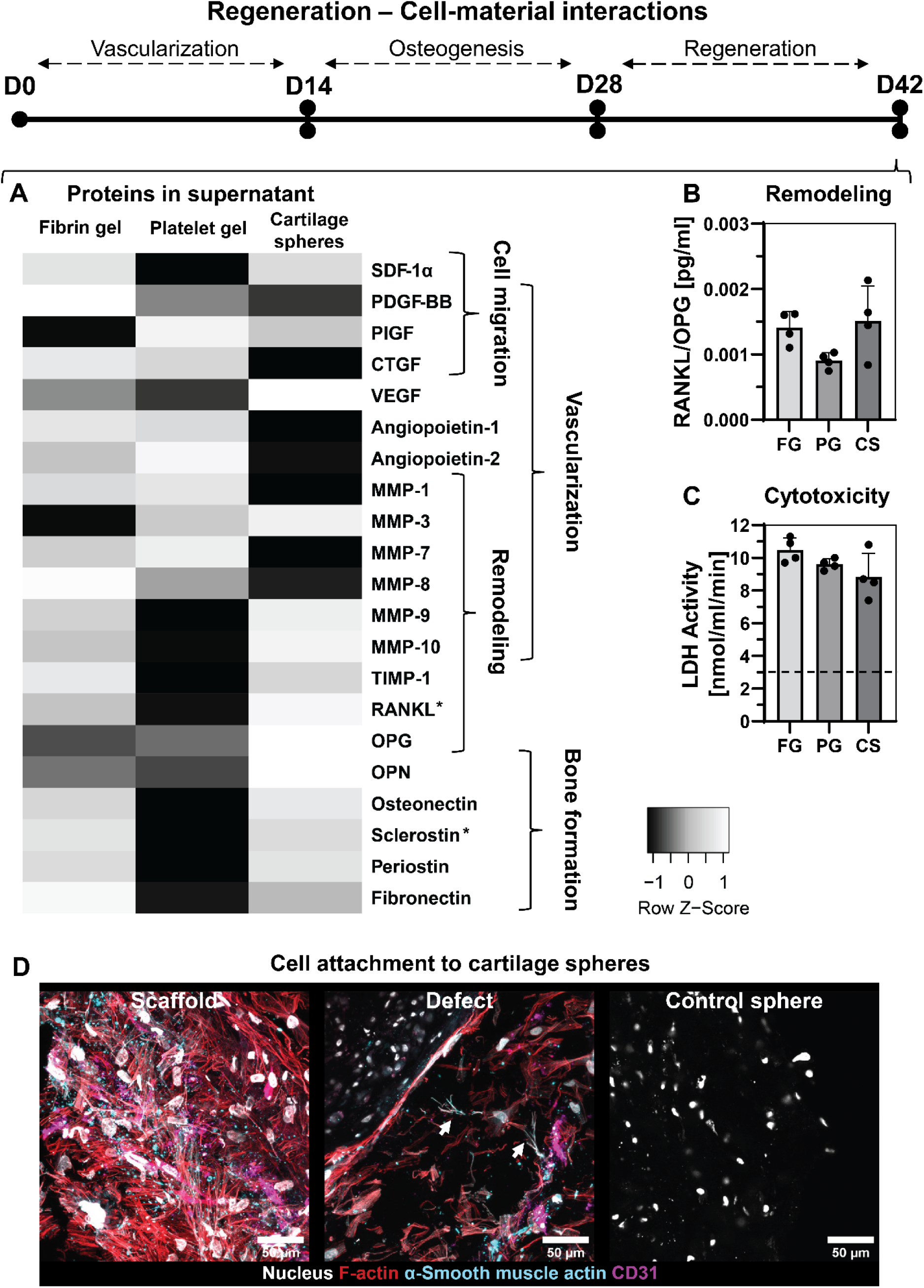
Evaluation of cell-material interactions after 42 days of culture (14 days post-implantation). (**A**) Multiplex immunoassays to measure proteins in cell supernatants, presented as z-scores (*i.e*., the normalized deviation from the average of all experimental groups). Concentration values were used for statistical comparisons, *p*<0.05 for RANKL and sclerostin (One-way ANOVA and Holm-Šídák’s post hoc tests), *ns* for other factors (Kruskal-Wallis tests for MMP-10, PlGF, MMP-3 and MMP-7, One-way ANOVAs for all other factors). (**B**) RANKL/OPG ratios in the culture medium supernatants, *ns* (One-way ANOVA). (**C**) Cytotoxicity measured by LDH activity in the supernatant. Dashed line represents concentration measured in control medium, *ns* (One-way ANOVA). (**D**) Z-stack maximum intensity projection images of sections stained for F-Actin (red), the nucleus (gray), endothelial cell marker CD31 (magenta) and supporting cell marker *α*-smooth muscle actin (cyan). White arrows point at locations where *α*-smooth muscle actin was co-localized with the actin cytoskeleton. Abbreviations: day (D), stromal derived factor (SDF), platelet derived growth factor (PDGF), placental growth factor (PlGF), connective tissue growth factor (CTGF), vascular endothelial growth factor (VEGF), matrix metalloproteinase (MMP), tissue inhibitor of metalloproteinase (TIMP), receptor activator of nuclear factor kappa-*β* ligand (RANKL), osteoprotegerin (OPG), osteopontin (OPN), lactate dehydrogenase (LDH). (The reader is referred to the web version of this article for colored images)

As such, proteins important for physiological bone regeneration can be captured with the presented bone defect model. Only sclerostin concentration in scaffolds implanted with platelet gel was at a similar level as the control medium (Table S2). In scaffolds implanted with fibrin gel and cartilage spheres, significantly more sclerostin was measured when comparing the concentrations of all experimental groups (Table S2). Sclerostin is known to inhibit bone formation and promote bone resorption by stimulating secretion of receptor activator of nuclear factor kappa-*β* ligand (RANKL) by osteocytes [40]. Indeed, RANKL was also significantly higher in scaffolds implanted with cartilage spheres than in scaffolds implanted with platelet gel (Table S2). Therefore, scaffolds implanted with cartilage spheres might be the most potent inducer of material degradation and remodeling. This was however not reflected in the ratio between RANKL and its inhibitor osteoprotegerin (OPG), which was similar among groups (Figure 5B).

Other protein levels of interest were the concentrations of connective tissue growth factor (CTGF), vascular endothelial growth factor (VEGF), matrix metalloproteinase (MMP)-3 and MMP-9. The CTGF concentration, important for cell condensation during regeneration [41], tended to be lowest in scaffolds of which the defects were implanted with cartilage spheres where cell condensation might not be necessary because of the implanted soft callus mimic. In contrary, the for vascularization important VEGF tended to have the highest concentration in scaffolds of which the defects were implanted with cartilage spheres. However, in the same condition the concentrations of other important factors for vascularization angiopoietin-1 and 2 tended to be relatively low. MMP-3, which is involved in the degradation of cartilage [42], and MMP-9, the in bone most abundant MMP also involved in endochondral ossification [43,44], tended to be relatively high in scaffolds of which defects were implanted with cartilage spheres. However, none of these concentrations differed significantly from the concentrations measured in the other conditions. LDH activity in the culture medium supernatant was measured as an indicator of material induced cytotoxicity. No significant differences were observed between the different materials (Figure 5C). Based on the attachment of HUVECs to the cartilage spheres, as observed in the cell migration evaluation (Figure 4F), the cell-material interaction was evaluated microscopically for this condition (Figure 5D). By staining for CD31 as an endothelial marker and *α*-smooth muscle actin as a mural cell marker, the co-effort of HUVECs and hBMSCs to induce cartilage sphere vascularization was studied. Both inside the bone defect (*i.e*., in the scaffold) as around the sphere, *α*-smooth muscle actin was located around HUVECs, identified with CD31 (Figure 5D). However, *α*-smooth muscle actin did not appear co-located with the actin cytoskeleton. As such, hBMSCs might have lost their supporting cell functionality upon osteogenic differentiation.

Only around the defect, cells were found with typical *α*-smooth muscle actin fibers, indicating that some hBMSCs were still capable of performing their supporting cell or pericyte-like cell functionality. In addition, by the absence of the endothelial marker CD31 around the spheres, also the attachment of hBMSCs to the spheres was confirmed by the layer of actin around them, which was not observed in non-implanted control spheres (Figure 5D). Infiltration of cells into the cartilage spheres was not observed.

### 3.4 Materials’ osteoinductive properties

After 42 days of culture, scaffolds and defects were evaluated for their cell and extracellular matrix content. In the scaffold, no clear differences between groups were observed in the DNA, hydroxyproline (*i.e*., a measure for collagen), and GAG content (Figure 6A-C). By measuring the DNA content in the defects, cell migration into the defect was quantified. In defects implanted with fibrin and platelet gel, little DNA was measured Figure 6D). Especially in defects implanted with platelet gel, DNA contents were low. In defects implanted with cartilage spheres, DNA content was much higher, likely caused by the presence of dead cartilage cells in the devitalized spheres. Interestingly, the measured DNA content in defects with implanted spheres tended to be lower than the measured DNA content in non-implanted control spheres. This suggests some degradation of implanted spheres. When visualizing collagen with picrosirius red, migrated cells in the defects filled with fibrin gel produced collagen (Figure 6E), which was characterized as collagen type I (Figure S3). This produced matrix did however show almost no mineralization (Figure 6I+K). Matrix formation and mineralization in the defect was not observed in defects implanted with platelet gel (Figure 6F+I+K). In defects implanted with cartilage spheres, collagen formation was observed around the spheres (Figure 6G).

**Figure 6.**
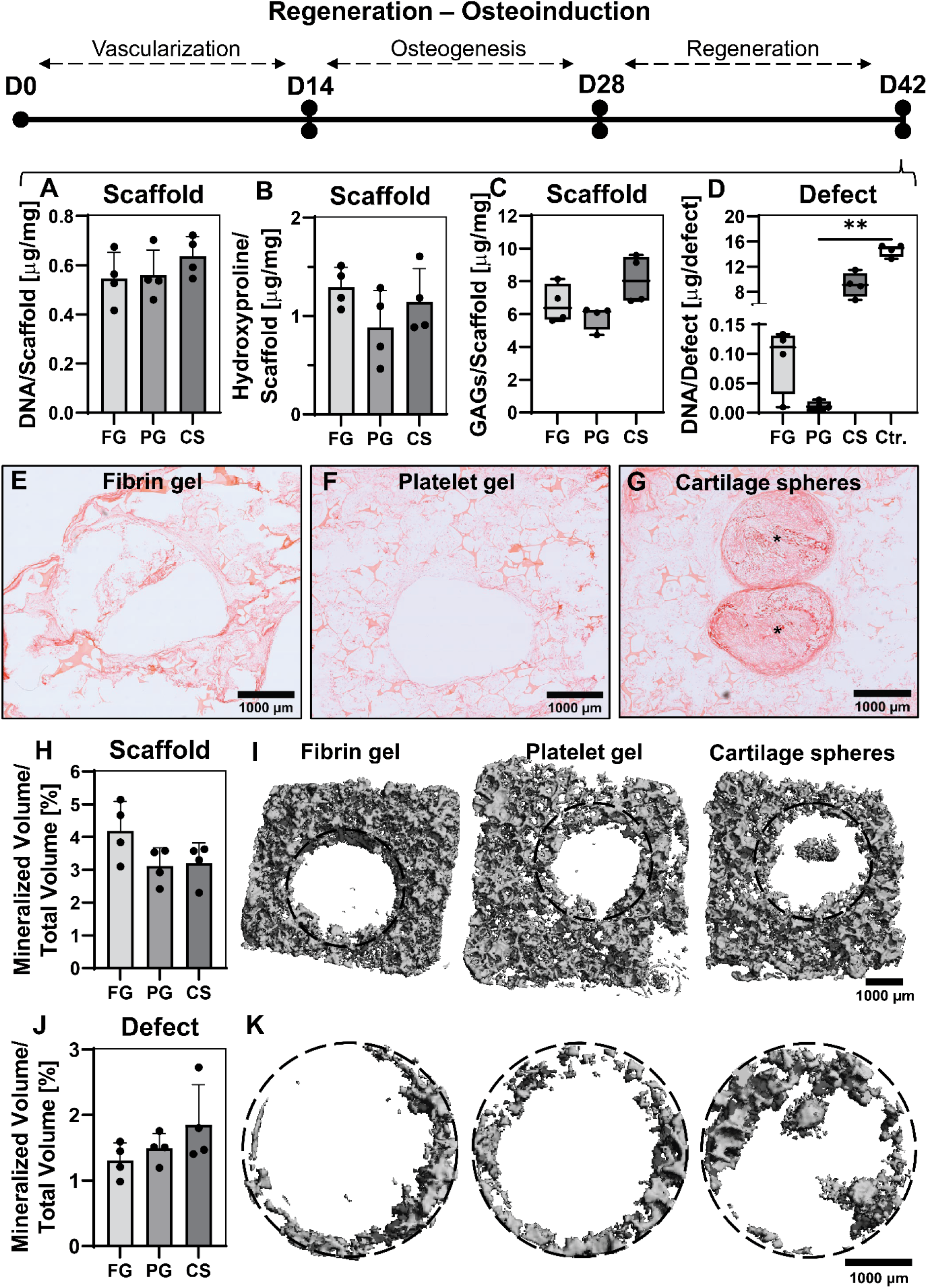
Evaluation of osteoinduction at the defect site after 42 days of culture (14 days post-implantation). (**A**) DNA quantification in scaffolds, ns (One-way ANOVA). (**B**) Hydroxyproline content quantification as a measure for collagen in scaffolds, ns (One-way ANOVA). (**C**) GAG content in scaffolds, ns (Kruskal-Wallis test). (**D**) DNA quantification in defects, *p*<0.05 (Kruskal-Wallis and Dunn’s post hoc tests, \*\**p*<0.01). (**E-G**) Collagen visualized by picrosirius red. Asterisks mark the cartilage spheres. (**H**) Quantified and (**I**) visualized mineralization of constructs using *µ*CT, *ns* (One-way ANOVA). (**J**) Quantified and (**K**) visualized mineralization of defects using *µ*CT, *ns* (One-way ANOVA). Abbreviations: day (D), glycosaminoglycan (GAG), micro-computed tomography (*µ*CT).

In addition, mineralization of the cartilage spheres was observed (Figure 6I+K). When quantifying the overall mineralization in the scaffold and the defects, no significant differences were observed. However, outer scaffolds implanted with fibrin gel tended to have a higher mineralized volume than outer scaffolds implanted with platelet gel or cartilage spheres, whereas defects implanted with cartilage spheres tended to have a higher mineralized volume compared to defects implanted with fibrin or platelet gel. This was caused by mineralization of some cartilage spheres. For model validation, cartilage sphere mineralization in our *in vitro* model was compared to cartilage sphere mineralization *in vivo* 14 days after ectopic implantation in a rat model. *In vivo*, cartilage spheres were also partly mineralized on day 14 (Figure S4).

## 4. Discussion

With the poor translation from *in vitro* assessments to *in vivo* models and clinical trials, there is a need for advanced *in vitro* models. For evaluation of materials for *in situ* or material-driven bone regeneration, where intelligent biomaterials make use of the bone’s innate capacity to regenerate and remodel upon implantation, such *in vitro* models do not currently exist [14]. This not only hinders clinical translation, but also leaves a significant burden on animal experiments [8]. Novel bone defect models to evaluate material-driven bone regeneration and to address the replacement, reduction and refinement of animal experiments principle (3Rs) have been developed in the recent years. This includes *ex vivo* human bone explant cultures and multiple semi-orthotopic implantations of bovine bone explants in mice [39,45]. These models allow for the evaluation of cell-material interactions of multiple cell types in their native environment. However, the multicellular environment also complicates unraveling cell-material interactions. Thereby, keeping bone explants viable over time outside the living body remains a challenge [13,46]. Cell death in explants transplanted in a different host as described above may trigger a non-physiological regeneration/remodeling response. Instead, with the development of an *in vitro* bone defect model a bottom-up approach can be adopted. As such, the complexity of models can be adapted, depending on which cell-material interactions need to be elucidated. Here, we used a tissue engineering approach to create an *in vitro* human bone defect model that enabled the *in vitro* evaluation of the material’s potential to stimulate vascularization, cell migration and osteoinduction.

A co-culture of HUVECs and hBMSCs was used to create a bone-like extracellular matrix with vascular-like structures. By using a SF scaffold with two *in vitro* “critically sized” defects [20], 3D tissue growth was facilitated around the defects. HUVECs were stimulated to form vascular-like structures in endothelial growth medium, using hBMSCs as supporting and osteogenic cells. These vascular-like structures indeed developed over a period of 14 days, and showed a branching morphology towards the end of the vascularization phase (i.e., from day 0 to day 14). Vascular-like structures were mostly maintained for an additional 14 days when osteogenic differentiation and bone-like matrix formation of hBMSCs was induced. Although the main focus was on the defect during the regeneration phase (i.e., from day 28 to day 42), some HUVEC-hBMSC networks had regressed at this stage, based on the fragmented α-smooth muscle actin staining as a supporting cell marker. In addition, one limitation of the presented study is the lack of quality assessment of the vascular network [47]. Therefore, future experiments might further investigate functional properties like network connectivity and whether the vascular-like structures are indeed hollow and maybe even perfusable. Additionally, vasculature in physiological bone has, based on its location and presumably function, a distinct molecular identity (i.e., a relative expression level of CD31 and endomucin) [48]. Bone-specific vasculature markers were not assessed in our study. Moreover, during physiological bone regeneration, vascularization is also a result of angiogenesis from existing vasculature [49], while in our *in vitro* model HUVECs only migrated from likely immature networks in the scaffold. Nevertheless, a primitive endothelial network, osteogenic differentiation and bone-like matrix formation, including the formation of collagen type I, osteopontin, and hydroxyapatite, was established prior to the implantation of the biomaterials.

To improve the translation from *in vitro* assessments to *in vivo* models and clinical trials, standardized protocols for the analyses of treatments on all model levels should be implemented [50]. This would allow comparison and potential extrapolation of experimental outcomes from different models [51]. Established biomarkers could facilitate in these comparisons [50,52,53]. For bone regeneration, commonly assessed biomarkers include *e.g*., alkaline phosphatase, RUNX2, bone morphogenetic protein 2 and 7, osteopontin, osteocalcin, collagen type I, and vascular markers VEGF and CD31 [54]. As bone regeneration also involves callus and woven bone remodeling, the regulators of bone turnover RANKL and OPG are relevant as well [54,55]. In our effort to improve and allow *in vitro/in vivo* translation, some of these markers were evaluated. In addition to these markers, other factors that have been reported to be involved in cell migration, vascularization, remodeling, and bone formation were evaluated upon material implantation (Table S2). An apparent difference was found in markers representative for the inhibition of bone formation and the stimulation of bone resorption; sclerostin and RANKL, respectively [55]. In scaffolds implanted with cartilage spheres, a higher level of sclerostin and RANKL was measured when compared to scaffolds implanted with platelet gel. *In vivo*, osteocytes can express sclerostin and RANKL in the absence of mechanical stimulation to regulate bone remodeling [56]. Scaffolds that expressed sclerostin and RANKL might have been more matured. In these scaffolds, hBMSCs might have been embedded in their matrix and differentiated into osteocytes, which were stimulated to produce sclerostin and RANKL in the absence of mechanical loading. However, this needs to be confirmed by the evaluation of morphological osteocyte characteristics and the localization of osteocyte markers [18].

While no clear differences in other biomarkers were observed, the presence of all markers in the culture medium supernatants allows for the evaluation of physiological processes in bone regeneration with the presented *in vitro* model. Most likely, measured protein concentrations represent the bulk expression from cells in the scaffold, rather than the expression of the limited number of cells that interacted with the materials. Nevertheless, our *in vitro* model was able to capture some physiological regeneration events. First, the chemotaxis needed to attract vasculature to the defect site was reflected by the model to some extent. While in the fibrin gel mostly hBMSCs migrated to the defect site, in the platelet gel and cartilage spheres conditions, also HUVECs migrated to the defect site. Second, *in vivo* bone regeneration includes soft callus mineralization [57], something that was also observed in our *in vitro* model. As such, the *in vitro* model presented in this study shows resemblance to some stages of physiological bone regeneration.

When comparing the obtained *in vitro* results to *in vivo* data, some similarities can be observed. It is well accepted that for *in vivo* bone defects, fibrin alone is incapable of inducing full regeneration [58,59]. This was also observed in the present *in vitro* model which might be explained by its inability to attract vascularization. Further, recently, blood clots were used to regenerate mouse cranial defects [60]. It was found that *in vivo* these blood clots were completely degraded 15 days after implantation [60]. In parallel, after 14 days, blood clot mimics in the current study were mostly degraded even though in our study a blood clot mimic was used instead of a physiological blood clot. Furthermore, in this study, the implanted devitalized cartilage spheres showed initial mineralization after 14 days, similar to when implanted *in vivo* for 14 days. Previously, these devitalized cartilage spheres were already shown to stimulate subcutaneous endochondral bone formation and bone regeneration in critically sized long bone defects in rats [29]. In these *in vivo* defects, complete bridging of the gap was already observed with *µ*CT 4 weeks after implantation [29]. The successful mineralization seen *in vivo* and *in vitro* might be attributed to the presence of alkaline phosphatase in the spheres [29]. However, cartilage spheres that were further differentiated into hypertrophic cartilage spheres and subsequently devitalized, containing higher levels of alkaline phosphatase, were less successful in defect regeneration *in vivo* [29]. Thus, while *in vitro* mineralization might be mainly caused by the presence of alkaline phosphatase, *in vivo* there are likely a multitude of factors regulating mineralization and regeneration. While some similarities were observed, the major difference between *in vivo* and *in vitro* models is the lack of the initial inflammatory response in *in vitro* models. Initial immune responses *in vivo* can be predictable for subsequent bone regeneration [61], which underlines the importance of the immune cells in bone regeneration. The immune system is highly complicated, featuring a multitude of cell types and interactions. Therefore, future studies should investigate which immune responses are predictive for bone regeneration and how these immune responses can be integrated into the *in vitro* model. The addition of monocytes and their subsequent macrophage and osteoclast differentiation might already allow for enhanced degradation of the materials which may, in case of the spheres, improve growth factor release from the spheres and vascular infiltration into the spheres.

Besides the lack of immune cells (*e.g*., monocytes, macrophages and osteoclasts), vascular maturation and functionality in the presented *in vitro* model, the model also lacks the presence of adjacent tissues that influence bone regeneration *in vivo* (*e.g*., periosteum, bone marrow, muscle tissue [62,63]. Animal experiments are therefore still inevitable. Other limitations are the absence of mechanical loading and the presence of the xenogeneic FBS. Mechanical loading is a well-accepted regulator of bone remodeling [64], and *in vivo* mechanical loading was also shown to influence bone regeneration [65]. Bioreactors could facilitate in this mechanical loading, which ideally allows for longitudinal microscopic evaluation and in which samples are easily accessible for staining and material implantation. To replace FBS, alternatives for HUVEC-hBMSC co-cultures like human platelet lysate or defined serum-free media need to be explored. Human platelet lysate was already demonstrated to support bone remodeling [66], while serum-free medium proved efficient for HUVECs co-cultured with human adipose tissue derived stromal cells [67].

## 5. Conclusion

Advanced *in vitro* human bone defect models could facilitate in the evaluation of materials for *in situ* bone regeneration, addressing both translational and ethical concerns regarding animal experiments. Here, we present such an *in vitro* model, which was used to implant physiologically relevant materials for bone regeneration including a fibrin gel, platelet gel as blood clot mimic, and cartilage spheres as soft callus mimics. Within this model, important hallmarks of *in situ* bone regeneration including cell migration, vascularization, and osteoinduction, were observer *in vitro*. These included the endothelial cell chemotaxis induced by the blood clot mimic and the mineralization of the soft callus mimic. As such, this *in vitro* model could contribute to improved pre-clinical evaluation while aiding to reduce the need for animal experiments.

## 6. Competing interest

The authors declare no competing interest.

## 7. Author Contributions

BdW, EC, KI, DG and SH contributed to conception, methodology and design of the study. BdW and EC performed the experiments and analysed the experimental results. Cartilage spheres were provided by LdS and DG. BdW wrote the original draft of the manuscript. All authors contributed to manuscript revision and approved the submitted version. BdW and SH acquired funding for this research.

## 8. Funding

This work is part of the research program TTW with project number TTW 016.Vidi.188.021, which is (partly) financed by the Netherlands Organization for Scientific Research (NWO). This research was also financially supported by the Gravitation Program “Materials Driven Regeneration”, funded by the Netherlands Organization for Scientific Research (024.003.013).

## 9. Acknowledgement

We thank Dewy van der Valk for the transport of the cartilage spheres from Utrecht to Eindhoven. We thank the Multiplex Core Facility of the Laboratory for Translational Immunology of the University Medical Center Utrecht, the Netherlands, for the Luminex analysis.

## Notes

### Competing Interest Statement

The authors have declared no competing interest.

## References

[1] R. Zura, Z. Xiong, T. Einhorn, J.T. Watson, R.F. Ostrum, M.J. Prayson, G.J. Della Rocca, S. Mehta, T. McKinley, Z. Wang, R.G. Steen, Epidemiology of Fracture Nonunion in 18 Human Bones, JAMA Surg. 151 (2016) e162775. https://doi.org/10.1001/jamasurg.2016.2775.

[2] L.A. Mills, S.A. Aitken, A.H.R.W. Simpson, The risk of non-union per fracture: current myths and revised figures from a population of over 4 million adults, Acta Orthop. 88 (2017) 434–439. https://doi.org/10.1080/17453674.2017.1321351.

[3] A.I.P.M. Smits, C.V.C. Bouten, Tissue Engineering meets Immunoengineering: Prospective on Personalized In Situ Tissue Engineering Strategies, Curr Opin Biomed Eng. 6 (2018) 17–26. https://doi.org/10.1016/j.cobme.2018.02.006.

[4] S. Cao, Y. Zhao, Y. Hu, L. Zou, J. Chen, New perspectives: In-situ tissue engineering for bone repair scaffold, Compos Part B. 202 (2020) 108445. https://doi.org/10.1016/j.compositesb.2020.108445.

[5] K.L. Spiller, S. Nassiri, C.E. Witherel, R.R. Anfang, J. Ng, K.R. Nakazawa, T. Yu, G. Vunjak-Novakovic, Sequential delivery of immunomodulatory cytokines to facilitate the M1-to-M2 transition of macrophages and enhance vascularization of bone scaffolds, Biomaterials. 37 (2015) 194–207. https://doi.org/10.1016/j.biomaterials.2014.10.017.

[6] T. Sun, C. Meng, Q. Ding, K. Yu, X. Zhang, W. Zhang, W. Tian, Q. Zhang, X. Guo, B. Wu, Z. Xiong, In situ bone regeneration with sequential delivery of aptamer and BMP2 from an ECM-based scaffold fabricated by cryogenic free-form extrusion, Bioact Mater. 6 (2021) 4163–4175. https://doi.org/10.1016/j.bioactmat.2021.04.013.

[7] O. Omar, T. Engstrand, L. Kihlström, B. Linder, J. Åberg, F.A. Shah, A. Palmquist, U. Birgersson, I. Elgali, M. Pujari-Palmer, H. Engqvist, P. Thomsen, In situ bone regeneration of large cranial defects using synthetic ceramic implants with a tailored composition and design, PNAS. 117 (2020) 26660–26671. https://doi.org/10.1073/pnas.2007635117.

[8] G. Hulsart-Billström, J.I. Dawson, S. Hofmann, R. Müller, M.J. Stoddart, M. Alini, H. Redl, A. El Haj, R. Brown, V. Salih, J. Hilborn, S. Larsson, R.O.C. Oreffo, A surprisingly poor correlation between in vitro and in vivo testing of biomaterials for bone regeneration: Results of a multicentre analysis, Eur Cells Mater. 31 (2016) 312–322. https://doi.org/10.22203/eCM.v031a20.

[9] D.W. Thomas, J. Burns, J. Audette, Adam Carroll, C. Dow-Hygelund, M. Hay, Clinical Development Success Rates 2006-2015, 2016.

[10] D. Thomas, D. Chancellor, A. Micklus, S. LaFever, M. Hay, S. Chaudhuri, R. Bowden, A.W. Lo, Clinical Development Success Rates and Contributing Factors 2011-2020, 2021.

[11] F. Barré-Sinoussi, X. Montagutelli, Animal models are essential to biological research: issues and perspectives, Futur Sci OA. 1 (2015) FSO63. https://doi.org/10.4155/FSO.15.63.

[12] A. Holmes, R. Brown, K. Shakesheff, Engineering tissue alternatives to animals: applying tissue engineering to basic research and safety testing, Regen Med. 4 (2009) 579–592. https://doi.org/10.2217/rme.09.26.

[13] E.E.A. Cramer, K. Ito, S. Hofmann, Ex vivo Bone Models and Their Potential in Preclinical Evaluation, Curr Osteoporos Rep. 19 (2021) 75–87. https://doi.org/10.1007/s11914-020-00649-5.

[14] S. Chen, X. Chen, Z. Geng, J. Su, The horizon of bone organoid: A perspective on construction and application, Bioact Mater. 18 (2022) 15–25. https://doi.org/10.1016/j.bioactmat.2022.01.048.

[15] S. Caddeo, M. Boffito, S. Sartori, Tissue engineering approaches in the design of healthy and pathological in vitro tissue models, Front Bioeng Biotechnol. 5 (2017) 40. https://doi.org/10.3389/fbioe.2017.00040.

[16] A.E. Gilchrist, B.A.C. Harley, Engineered Tissue Models to Replicate Dynamic Interactions within the Hematopoietic Stem Cell Niche, Adv Healthc Mater. 11 (2022) 2102130. https://doi.org/10.1002/adhm.202102130.

[17] Q. Xiong, N. Zhang, M. Zhang, M. Wang, L. Wang, Y. Fan, C.Y. Lin, Engineer a pre-metastatic niched microenvironment to attract breast cancer cells by utilizing a 3D printed polycaprolactone/nano-hydroxyapatite osteogenic scaffold –An in vitro model system for proof of concept, J Biomed Mater Res -Part B Appl Biomater. 110 (2022) 1604–1614. https://doi.org/10.1002/jbm.b.35021.

[18] A. Akiva, J. Melke, S. Ansari, N. Liv, R. van der Meijden, M. van Erp, F. Zhao, M. Stout, W.H. Nijhuis, C. de Heus, C. Muñiz Ortera, J. Fermie, J. Klumperman, K. Ito, N. Sommerdijk, S. Hofmann, An Organoid for Woven Bone, Adv Funct Mater. 31 (2021) 2010524. https://doi.org/10.1002/adfm.202010524.

[19] S.J.A. Remmers, B.W.M. de Wildt, M.A.M. Vis, E.S.R. Spaander, R.B.M. de Vries, K. Ito, S. Hofmann, Osteoblast-osteoclast co-cultures: A systematic review and map of available literature, PLoS One. 16 (2021) e0257724. https://doi.org/10.1371/journal.pone.0257724.

[20] J.R. Vetsch, R. Müller, S. Hofmann, The influence of curvature on threedimensional mineralized matrix formation under static and perfused conditions: An in vitro bioreactor model, J R Soc Interface. 13 (2016) 20160425. https://doi.org/10.1098/rsif.2016.0425.

[21] G. Matziolis, J. Tuischer, G. Kasper, M. Thompson, B. Bartmeyer, D. Krocker, C. Perka, G. Duda, Simulation of Cell Differentiation in Fracture Healing: Mechanically Loaded Composite Scaffolds in a Novel Bioreactor System, Tissue Eng. 12 (2006) 201–208. https://doi.org/10.1089/ten.2006.12.201.

[22] K.D. Hankenson, M. Dishowitz, C. Gray, M. Schenker, Angiogenesis in bone regeneration, Injury. 42 (2011) 556–561. https://doi.org/10.1016/j.injury.2011.03.035.

[23] S. Verrier, M. Alini, E. Alsberg, S.R. Buchman, D. Kelly, M.W. Laschke, M.D. Menger, W.L. Murphy, J.P. Stegemann, M. Schütz, T. Miclau, M.J. Stoddart, C. Evans, Tissue engineering and regenerative approaches to improving the healing of large bone defects, Eur Cells Mater. 32 (2016) 87–110. https://doi.org/10.22203/eCM.v032a06.

[24] S. Shafiee, S. Shariatzadeh, A. Zafari, A. Majd, H. Niknejad, Recent Advances on Cell-Based Co-Culture Strategies for Prevascularization in Tissue Engineering, Front Bioeng Biotechnol. 9 (2021) 745314. https://doi.org/10.3389/fbioe.2021.745314.

[25] K. Pill, S. Hofmann, H. Redl, W. Holnthoner, Vascularization mediated by mesenchymal stem cells from bone marrow and adipose tissue: A comparison, Cell Regen. 4 (2015) 8. https://doi.org/10.1186/s13619-015-0025-8.

[26] A. Grosso, M.G. Burger, A. Lunger, D.J. Schaefer, A. Banfi, N. Di Maggio, It takes two to tango: Coupling of angiogenesis and osteogenesis for bone regeneration, Front Bioeng Biotechnol. 5 (2017) 68. https://doi.org/10.3389/fbioe.2017.00068.

[27] I. Pennings, L.A. van Dijk, J. van Huuksloot, J.O. Fledderus, K. Schepers, A.K. Braat, E.C. Hsiao, E. Barruet, B.M. Morales, M.C. Verhaar, A.J.W.P. Rosenberg, D. Gawlitta, Effect of donor variation on osteogenesis and vasculogenesis in hydrogel cocultures, J Tissue Eng Regen Med. 13 (2019) 433–445. https://doi.org/10.1002/term.2807.

[28] B.J. Klotz, K.S. Lim, Y.X. Chang, B.G. Soliman, I. Pennings, F.P.W. Melchels, T.B.F. Woodfield, A.J.W.P. Rosenberg, J. Malda, D. Gawlitta, Engineering of a complex bone tissue model with endothelialised channels and capillary-like networks, Eur Cells Mater. 35 (2018) 335–349. https://doi.org/10.22203/eCM.v035a23.

[29] A. Longoni, L. Utomo, A. Robinson, R. Levato, A.J.W.P. Rosenberg, D. Gawlitta, Acceleration of Bone Regeneration Induced by a Soft-Callus Mimetic Material, Adv Sci. 9 (2022) 2103284. https://doi.org/10.1002/advs.202103284.

[30] S. Hofmann, H. Hagenmüller, A.M. Koch, R. Müller, G. Vunjak-Novakovic, D.L. Kaplan, H.P. Merkle, L. Meinel, Control of in vitro tissue-engineered bone-like structures using human mesenchymal stem cells and porous silk scaffolds, Biomaterials. 28 (2007) 1152–1162. https://doi.org/10.1016/j.biomaterials.2006.10.019.

[31] J. Melke, F. Zhao, K. Ito, S. Hofmann, Orbital seeding of mesenchymal stromal cells increases osteogenic differentiation and bone -like tissue formation, J Orthop Res. 2020 (2019) 1–10. https://doi.org/10.1002/jor.24583.

[32] S.J.A. Aper, A.C.C. Van Spreeuwel, M.C. Van Turnhout, A.J. Van Der Linden, P.A. Pieters, N.L.L. Van Der Zon, S.L. De La Rambelje, C.V.C. Bouten, M. Merkx, Colorful protein-based fluorescent probes for collagen imaging, PLoS One. 9 (2014) e114983. https://doi.org/10.1371/journal.pone.0114983.

[33] J. Ge, D.K. Wood, D.M. Weingeist, S. Prasongtanakij, P. Navasumrit, M. Ruchirawat, B.P. Engelward, Standard fluorescent imaging of live cells is highly genotoxic, Cytom Part A. 83 (2013) 552–560. https://doi.org/10.1002/cyto.a.22291.

[34] M.L. Bouxsein, S.K. Boyd, B.A. Christiansen, R.E. Guldberg, K.J. Jepsen, R. Müller, Guidelines for assessment of bone microstructure in rodents using micro-computed tomography, J Bone Miner Res. 25 (2010) 1468–1486. https://doi.org/10.1002/jbmr.141.

[35] G. Huszar, J. Maiocco, F. Naftolin, Monitoring of collagen and collagen fragments in chromatography of protein mixtures, Anal Biochem. 105 (1980) 424–429. https://doi.org/10.1016/0003-2697(80)90481-9.

[36] R. Farndale, C. Sayers, A. Barrett, A direct spectrophotometric microassay for sulfated glycosaminoglycans in cartilage cultures., Connect Tissue Res. 9 (1982) 247–248.

[37] J. Schindelin, I. Arganda-Carreras, E. Frise, V. Kaynig, M. Longair, T. Pietzsch, S. Preibisch, C. Rueden, S. Saalfeld, B. Schmid, J.Y. Tinevez, D.J. White, V. Hartenstein, K. Eliceiri, P. Tomancak, A. Cardona, Fiji: An open-source platform for biological-image analysis, Nat Methods. 9 (2012) 676–682. https://doi.org/10.1038/nmeth.2019.

[38] A. Rutkovskiy, K.-O. Stensløkken, I.J. Vaage, Osteoblast Differentiation at a Glance, Med Sci Monit Basic Res. 22 (2016) 95–106. https://doi.org/10.12659/MSMBR.901142.

[39] T. Klüter, R. Hassan, A. Rasch, H. Naujokat, F. Wang, P. Behrendt, S. Lippross, L. Gerdesmeyer, D. Eglin, A. Seekamp, S. Fuchs, An ex vivo bone defect model to evaluate bone substitutes and associated bone regeneration processes, Tissue Eng Part C Methods. 26 (2020). https://doi.org/10.1089/ten.TEC.2019.0274.

[40] P.K. Suen, L. Qin, Sclerostin, an emerging therapeutic target for treating osteoporosis and osteoporotic fracture: A general review, J Orthop Transl. 4 (2016) 1–13. https://doi.org/10.1016/j.jot.2015.08.004.

[41] J.A. Arnott, A.G. Lambi, C.M. Mundy, H. Hendesi, R.A. Pixley, T.A. Owen, F.F. Safadi, S.N. Popoff, The Role of Connective Tissue Growth Factor (CTGF/CCN2) in Skeletogenesis, Crit Rev Eukaryot Gene Expr. 21 (2011) 43–69. https://doi.org/10.1615/critreveukargeneexpr.v21.i1.40.

[42] J. Wan, G. Zhang, X. Li, X. Qiu, J. Ouyang, J. Dai, S. Min, Matrix Metalloproteinase 3: A Promoting and Destabilizing Factor in the Pathogenesis of Disease and Cell Differentiation, Front Physiol. 12 (2021) 663978. https://doi.org/10.3389/fphys.2021.663978.

[43] K.B.S. Paiva, J.M. Granjeiro, Matrix Metalloproteinases in Bone Resorption, Remodeling, and Repair, in: Prog Mol Biol Transl Sci, Elsevier B.V., 2017: pp. 203–303. https://doi.org/10.1016/bs.pmbts.2017.05.001.

[44] N. Ortega, D.J. Behonick, Z. Werb, Matrix remodeling during endochondral ossification, Trends Cell Biol. 14 (2004) 86–93. https://doi.org/10.1016/j.tcb.2003.12.003.

[45] E. Andrés Sastre, Y. Nossin, I. Jansen, N. Kops, C. Intini, J. Witte-Bouma, B. van Rietbergen, S. Hofmann, Y. Ridwan, J.P. Gleeson, F.J. O’Brien, E.B. Wolvius, G.J.V.M. van Osch, E. Farrell, A new semi-orthotopic bone defect model for cell and biomaterial testing in regenerative medicine, Biomaterials. 279 (2021) 121187. https://doi.org/10.1016/j.biomaterials.2021.121187.

[46] T. Bellido, J. Delgado-Calle, Ex Vivo Organ Cultures as Models to Study Bone Biology, JBMR Plus. 4 (2020) e10345. https://doi.org/10.1002/jbm4.10345.

[47] X. Meng, Y. Xing, J. Li, C. Deng, Y. Li, X. Ren, D. Zhang, Rebuilding the Vascular Network: In vivo and in vitro Approaches, Front Cell Dev Biol. 9 (2021) 639299. https://doi.org/10.3389/fcell.2021.639299.

[48] E.C. Watson, R.H. Adams, Biology of bone: The vasculature of the skeletal system, Cold Spring Harb Perspect Med. 8 (2018) a031559. https://doi.org/10.1101/cshperspect.a031559.

[49] S. Stegen, N. van Gastel, G. Carmeliet, Bringing new life to damaged bone: The importance of angiogenesis in bone repair and regeneration, Bone. 70 (2015) 19–27. https://doi.org/10.1016/j.bone.2014.09.017.

[50] M. Wehling, Translational medicine: Can it really facilitate the transition of research “from bench to bedside”?, Eur J Clin Pharmacol. 62 (2006) 91–95. https://doi.org/10.1007/s00228-005-0060-4.

[51] W.B. Mattes, In vitro to in vivo translation, Curr Opin Toxicol. 23–24 (2020) 114–118. https://doi.org/10.1016/j.cotox.2020.09.001.

[52] A. Wendler, M. Wehling, Translatability scoring in drug development: Eight case studies, J Transl Med. 10 (2012) 39. https://doi.org/10.1186/1479-5876-10-39.

[53] J. Yadav, M. El Hassani, J. Sodhi, V.M. Lauschke, J.H. Hartman, L.E. Russell, Recent developments in in vitro and in vivo models for improved translation of preclinical pharmacokinetics and pharmacodynamics data, Drug Metab Rev. 53 (2021) 207–233. https://doi.org/10.1080/03602532.2021.1922435.

[54] S. Albeshri, A. Alblaihess, A.A. Niazy, S. Ramalingam, C. Sundar, H.S. Alghamdi, Biomarkers as independent predictors of bone regeneration around biomaterials: A systematic review of literature, J Contemp Dent Pract. 19 (2018) 605–6 18. https://doi.org/10.5005/jp-journals-10024-2306.

[55] T.R. Kuo, C.H. Chen, Bone biomarker for the clinical assessment of osteoporosis: Recent developments and future perspectives, Biomark Res. 5 (2017) 18. https://doi.org/10.1186/S40364-017-0097-4.

[56] L. Wang, X. You, L. Zhang, C. Zhang, W. Zou, Mechanical regulation of bone remodeling, Bone Res. 10 (2022) 16. https://doi.org/10.1038/s41413-022-00190-4.

[57] M. Ansari, Bone tissue regeneration: biology, strategies and interface studies, Prog Biomater. 8 (2019) 223–237. https://doi.org/10.1007/s40204-019-00125-z.

[58] A. Noori, S.J. Ashrafi, R. Vaez-Ghaemi, A. Hatamian-Zaremi, T.J. Webster, A review of fibrin and fibrin composites for bone tissue engineering, Int J Nanomedicine. 12 (2017) 4937–4961. https://doi.org/10.2147/IJN.S124671.

[59] M.K.E. Koolen, A. Longoni, J. Van Der Stok, O. Van Der Jagt, D. Gawlitta, H. Weinans, Complete regeneration of large bone defects in rats with commercially available fibrin loaded with BMP-2, Eur Cells Mater. 38 (2019) 94–105. https://doi.org/10.22203/eCM.v038a08.

[60] Q. Fan, J. Bai, H. Shan, Z. Fei, H. Chen, J. Xu, Q. Ma, X. Zhou, C. Wang, Implantable blood clot loaded with BMP-2 for regulation of osteoimmunology and enhancement of bone repair, Bioact Mater. 6 (2021) 4014–4026. https://doi.org/10.1016/j.bioactmat.2021.04.008.

[61] A. Cheng, C.E. Vantucci, L. Krishnan, M.A. Ruehle, T. Kotanchek, L.B. Wood, K. Roy, R.E. Guldberg, Early systemic immune biomarkers predict bone regeneration after trauma, PNAS. 118 (2021) e2017889118. https://doi.org/10.1073/pnas.2017889118.

[62] K. Shah, Z. Majeed, J. Jonason, R.J. O’Keefe, The role of muscle in bone repair: The cells, signals, and tissue responses to injury, Curr Osteoporos Rep. 11 (2013) 130–135. https://doi.org/10.1007/s11914-013-0146-3.

[63] T. Paul Neagu, M. Tiglis, I. Cocolos, C. Radu Jecan, The relationship between periosteum and fracture healing, Rom J Morphol Embryol. 57 (2016) 1215–1220.

[64] J. Wolff, The Law of Bone Remodelling, 1986.

[65] D.C. Betts, R. Müller, Mechanical regulation of bone regeneration: Theories, models, and experiments, Front Endocrinol (Lausanne). 5 (2014) 211. https://doi.org/10.3389/fendo.2014.00211.

[66] B.W.M. de Wildt, K. Ito, S. Hofmann, Human Platelet Lysate as Alternative of Fetal Bovine Serum for Enhanced Human In Vitro Bone Resorption and Remodeling, Front Immunol. 13 (2022) 915277. https://doi.org/10.3389/fimmu.2022.915277.

[67] B. Andrée, H. Ichanti, S. Kalies, A. Heisterkamp, S. Strauß, P.M. Vogt, A. Haverich, A. Hilfiker, Formation of three-dimensional tubular endothelial cell networks under defined serum-free cell culture conditions in human collagen hydrogels, Sci Rep. 9 (2019) 5437. https://doi.org/10.1038/s41598-019-41985-6.

