## Supporting information for "Evaluating material-driven regeneration in a tissue engineered human *in vitro* bone defect model"

The antibodies that were used for immunofluorescent stainings are listed in Table S1.

**Table S1.** List of antibodies that were used in this study.

| Antigen | Supplier | Catalogue No. | Conjugate | Species | Dilution |
|---|---|---|---|---|---|
| RUNX2 | Abcam | ab23981 | | Rabbit | 1:500 |
| Osteopontin | Thermo Fisher | 14-9096-82 | | Mouse | 1:200 |
| Collagen type 1 | Abcam | Ab34710 | | Rabbit | 1:200 |
| CD31 | Abcam | Ab37259 | | Mouse | 1:100 |
| α-Smooth muscle actin | Thermo Fisher | MA5-13188 | | Rabbit | 1:500 |
| Anti-rabbit IgG | Molecular Probes | A21246 | Alexa-647 | Goat | 1:200 |
| Anti-mouse IgG1 | Molecular Probes | A21127 | Alexa-555 | Goat | 1:200 |

Abbreviations: runt-related transcription factor 2 (RUNX2).

To study the influence of staining with OsteoSense™ 680, CNA35-mCherry and NucBlue™ Hoechst 33342 on cell death, unstained and stained samples were compared for their lactate dehydrogenase (LDH) release in the supernatant from day 40 to day 42 (Figure S1).

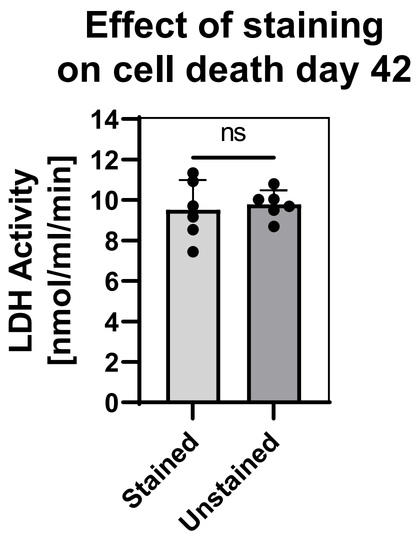


**Figure S1.** Potential staining-induced cell-death measured by LDH activity in the supernatant. Staining of samples did not induce additional cell death, *ns* (independent t-test).

On day 42 of the culture, cells had migrated to the implanted material in all conditions, but in defects implanted with fibrin gel, only hBMSCs migrated while in defects implanted with platelet gel both hBMSCs and HUVECs migrated (Figure S1). The migration of these cells appeared only in distinct areas. Defects implanted with cartilage spheres seemed to remain stable over time. In these defects, cell migration of both hBMSCs and HUVECs was observed by the presence of both cells around the spheres (Figure S1).

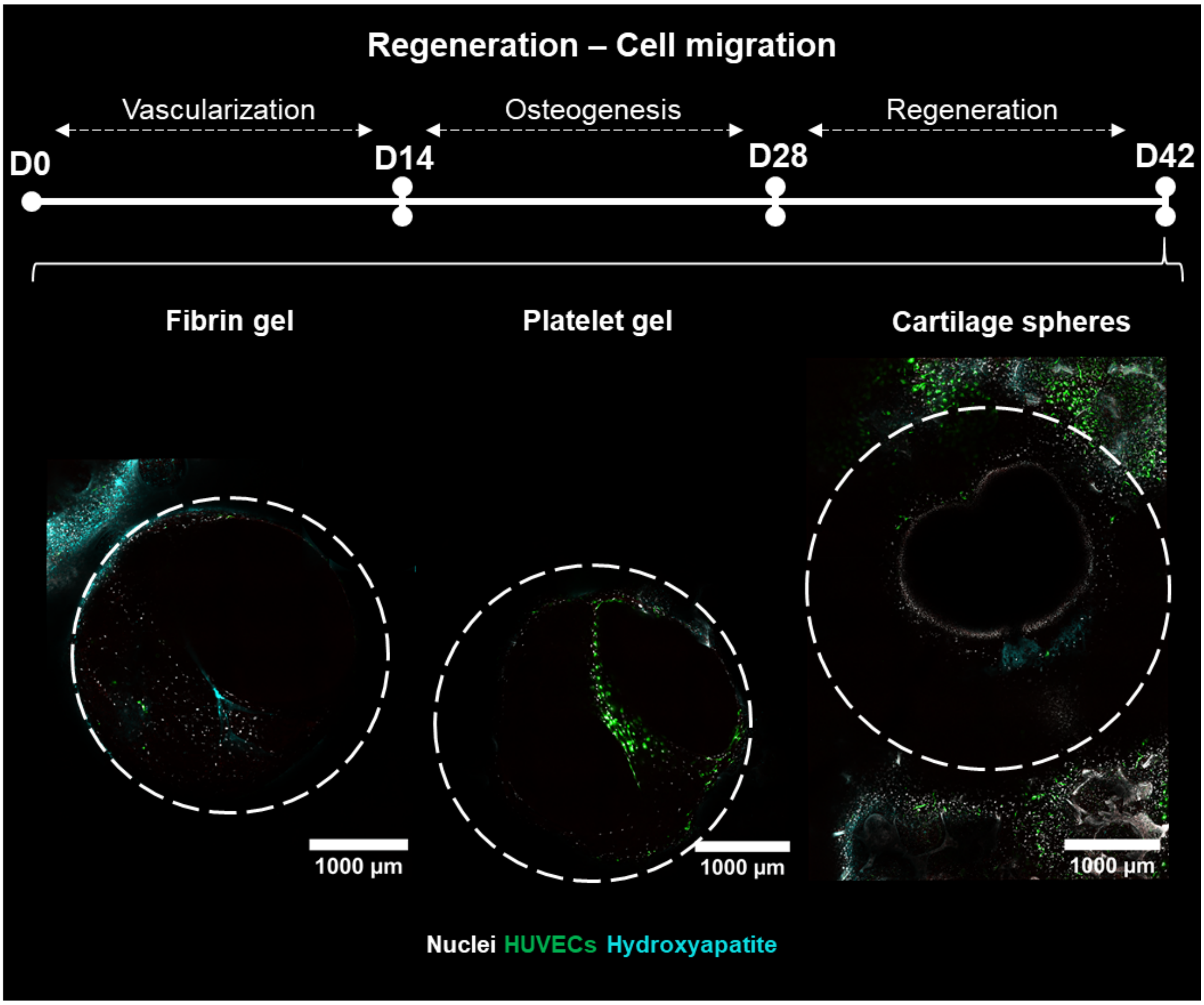


**Figure S2.** Evaluation of the regeneration phase of viable constructs. Images present defect overview images at day 42 of culture of constructs containing GFP-expressing HUVECs (green) and stained for nuclei (gray), and hydroxyapatite (cyan).

**Table S2.** Evaluation of proteins relevant in for bone regeneration using multiplex immunoassays.

| Analyte | Contribution to bone regeneration | Concentration in control medium (pg/ml) | Concentration in cell supernatant (pg/ml) |
|---|---|---|---|
| SDF-1α | Important role in migration of MSCs, mainly in inflammation [1]. Could support osteogenic differentiation and angiogenesis [2,3]. Hypothesized to recruit osteoclast precursors [4]. | 71.9 | **FG**: 372 ± 46.4 |
| | | | **PG**: 335 ± 74.0 |
| | | | **CS**: 370 ± 49.9 |
| PDGF-BB | Can enhance osteoclast differentiation of macrophage-like cells [5].<br>Overexpression of PDGF-BB in MSCs can promote osteogenesis and angiogenesis [6]. | 16.4 | **FG**: 43.5 ± 11.2 |
| | | | **PG**: 37.6 ± 4.75 |
| | | | **CS**: 35.8 ± 1.42 |
| PlGF° | Can recruit hematopoietic cells and stimulate the expression of pro-angiogenesis factors upon a fracture. Might influence osteogenic differentiation of MSCs and important for remodeling of healed fracture [7]. | 18.9 | **FG**: 764 ± 117 |
| | | | **PG**: 782 ± 161 |
| | | | **CS**: 787 ± 29.1 |
| CTGF | Important for MSC condensation and chondrogenic differentiation during endochondral bone regeneration [8]. | 14.0 | **FG**: 181 ± 28.7 |
| | | | **PG**: 179 ± 18.4 |
| | | | **CS**: 170 ± 23.5 |
| VEGF | Could enhance osteoclast survival and resorption [9,10]. Could also recruit chondroclasts, osteoclasts and immune cells. Promotes vascularization and can promote osteogenic differentiation [11]. | 47.9 | **FG**: $5.12*10^3 \pm 1.29*10^3$ |
| | | | **PG**: $4.83*10^3 \pm 544$ |
| | | | **CS**: $5.98*10^3 \pm 482$ |
| Angiopoietin-1 | Can promote vascular integrity in regenerating vasculature [12]. | 979 | **FG**: $10.5*10^3 \pm 1.58*10^3$ |
| | | | **PG**: $10.5*10^3 \pm 2.32*10^3$ |
| | | | **CS**: $9.91*10^3 \pm 1.21*10^3$ |
| Angiopoietin-2 | Can improve mineralization and angiogenesis in bone regeneration [13]. | $1.22*10^3$ | **FG**: $5.09*10^3 \pm 1.01*10^3$ |
| | | | **PG**: $5.72*10^3 \pm 1.08*10^3$ |
| | | | **CS**: $4.24*10^3 \pm 1.17*10^3$ |

| | | | |
|---|---|---|---|
| MMP-1 | Most abundant collagenase in tissues, cleaves collagen, but its absence only leads to modest abnormalities in bone remodeling [14]. | ND | **FG**: 785 ± 481<br>**PG**: 802 ± 401<br>**CS**: 655 ± 99.5 |
| MMP-3° | Involved in degradation of cartilage and the invasion of vasculature in osteoarthritis [15]. Can cleave non-collagenous proteins [14]. | 96.3 | **FG**: 294 ± 178<br>**PG**: 424 ± 376<br>**CS**: 454 ± 591 |
| MMP-7° | Can promote RANKL availability and cleave non-collagenous proteins. Might be crucial for proper bone regeneration [14]. | 445 | **FG**: $9.18*10^3 \pm 5.03*10^3$<br>**PG**: $10.4*10^3 \pm 9.63*10^3$<br>**CS**: $7.62*10^3 \pm 1.15*10^3$ |
| MMP-8 | Seems to be crucial for proper bone regeneration [14]. | 41.9 | **FG**: 155 ± 17.5<br>**PG**: 142 ± 19.6<br>**CS**: 133 ± 3.28 |
| MMP-9 | Most abundant MMP in bone, participates in osteoclast recruitment and the release of growth factors from the extracellular matrix [14]. MMP9 is also involved in endochondral ossification [16]. | 228 | **FG**: $1.04*10^3 \pm 78.7$<br>**PG**: 928 ± 125<br>**CS**: $1.08*10^3 \pm 33.1$ |
| MMP-10° | Might promote (pathological) calcification [17,18]. | 91.3 | **FG**: 529 ± 38.1<br>**PG**: 470 ± 22.2<br>**CS**: 558 ± 81.2 |
| TIMP-1 | High affinity for MMP-9 (inhibition), important for osteogenic lineage commitment of MSCs [14]. | 101 | **FG**: $18.9*10^3 \pm 448$<br>**PG**: $18.0*10^3 \pm 664$<br>**CS**: $18.8*10^3 \pm 621$ |
| RANKL | Typically expressed by cells from the osteogenic lineage, including MSCs, osteoblasts and osteocytes. Required for osteoclast differentiation [19]. | ND | **FG**: 19.6 ± 3.64<br>**PG**: 13.3 ± 3.80<br>**CS**: 24.1 ± 4.43 * |
| OPG | Can prevent RANKL from binding to the RANK receptor on | $1.41*10^3$ | **FG**: $14.2*10^3 \pm 3.56*10^3$<br>**PG**: $14.5*10^3 \pm 2.41*10^3$ |

| | | | | |
|---|---|---|---|---|
| | preosteoclasts, thereby inhibiting osteoclast differentiation [19,20]. | | **CS**: 16.9*$10^3$ ± 3.71*$10^3$ | |
| Osteopontin | Instrumental for intrafibrillar mineralization and promotes osteoclast activation [21].<br>Can promote osteoclast precursor migration [22]. | 321 | **FG**: 716 ± 96.7 | |
| | | | **PG**: 713 ± 15.1 | |
| | | | **CS**: 737 ± 45.7 | |
| Osteonectin | Can promote bone formation and mineralization. Regulates collagen fibrillogenesis [23]. | 1.27*$10^3$ | **FG**: 42.7*$10^3$ ± 5.55*$10^3$ | |
| | | | **PG**: 35.6*$10^3$ ± 5.69*$10^3$ | |
| | | | **CS**: 44.0*$10^3$ ± 4.20*$10^3$ | |
| Sclerostin | Inhibits bone formation and osteogenesis, could stimulate RANKL secretion by osteocytes, thereby promoting osteoclast differentiation [24]. | 414 | **FG**: 503 ± 36.5 | |
| | | | **PG**: 413 ± 41.0 | * (FG vs PG) |
| | | | **CS**: 496 ± 50.2 | * (PG vs CS) |
| Periostin | Upregulated in response to bone injury. Is crucial for callus, cartilage and bone formation [25]. | ND | **FG**: 61.0*$10^3$ ± 6.00*$10^3$ | |
| | | | **PG**: 56.8*$10^3$ ± 5.74*$10^3$ | |
| | | | **CS**: 61.3*$10^3$ ± 5.34*$10^3$ | |
| Fibronectin | Could inhibit osteoclastogenesis [26,27], but could enhance mature osteoclast activity and resorption [26].<br>Could promote osteogenic differentiation and bone-like matrix formation of MSCs at low coating densities, and inhibit differentiation but promote proliferation at higher coating densities [28]. | 330*$10^3$ | **FG**: 4.42*$10^6$ ± 621*$10^3$ | |
| | | | **PG**: 3.45*$10^6$ ± 456*$10^3$ | |
| | | | **CS**: 3.92*$10^6$ ± 758*$10^3$ | |

Values represent mean ± standard deviation or °median ± interquartile range. *$p$<0.05.

Abbreviations: stromal derived factor (SDF), platelet derived growth factor (PDGF), placental growth factor (PIGF), connective tissue growth factor (CTGF), vascular endothelial growth factor (VEGF), matrix metalloproteinase (MMP), tissue inhibitor of metalloproteinase (TIMP), receptor activator of nuclear factor kappa-$\beta$ ligand (RANKL), osteoprotegerin (OPG), osteopontin (OPN).

Collagen type I formation after 42 days was visualized in and around the defects using immunohistochemistry (Figure S2). Collagen type I formation was mainly visible in scaffolds implanted with fibrin gel or cartilage spheres (Figure S2).

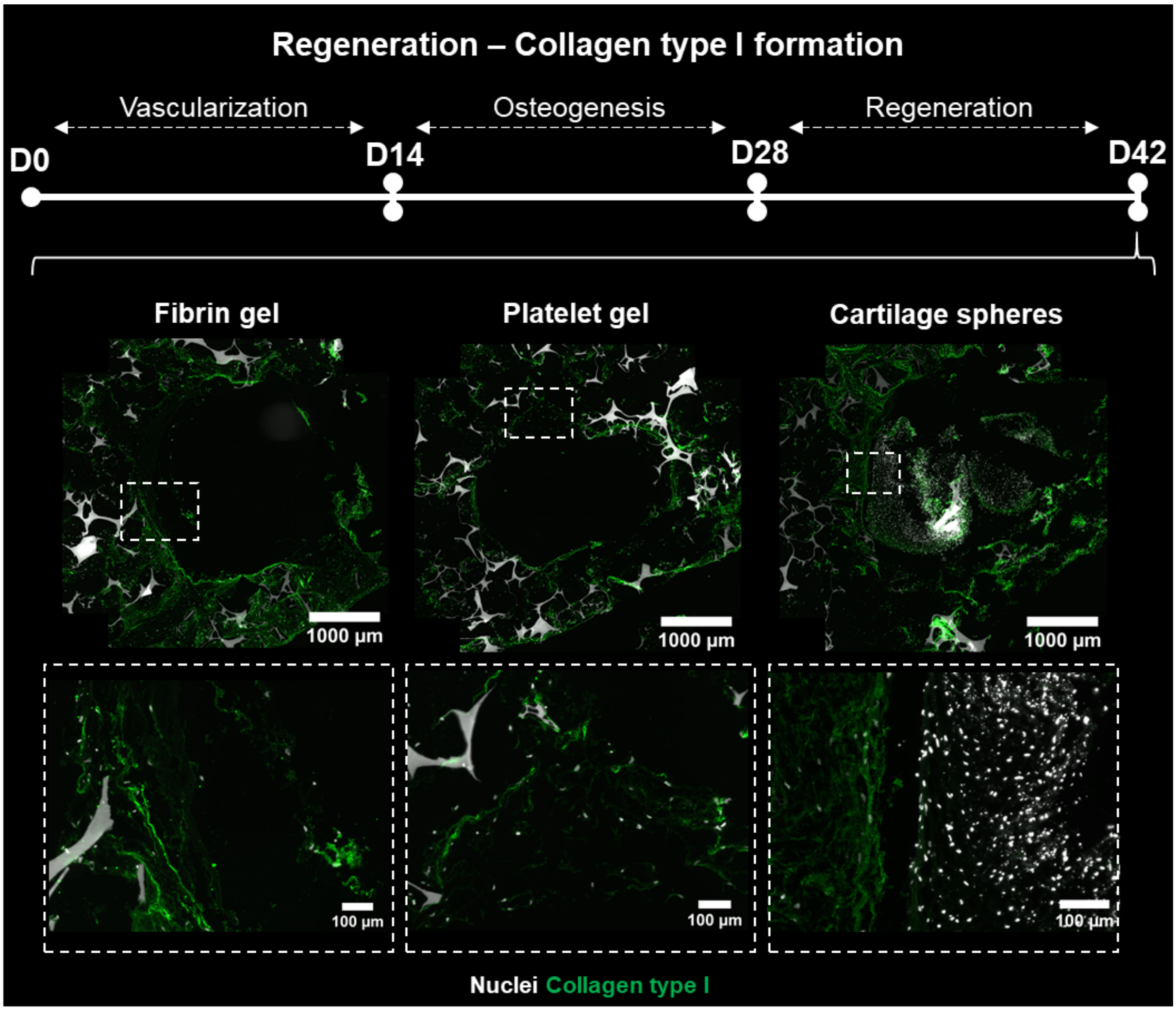


**Figure S3.** Collagen type I (green) immunohistochemical analysis. Top panel presents defect overview images, bottom panel present close-up images.

For model validation, *in vitro* cartilage sphere mineralization was compared to *in vivo* cartilage sphere mineralization 14 days after ectopic implantation in a rat model. *In vivo*, cartilage spheres were partly mineralized after 14 days, which is comparable to our *in vitro* model where parts of cartilage spheres were mineralized 14 days after artificial implantation.

**Mineralization D14 *in vivo***

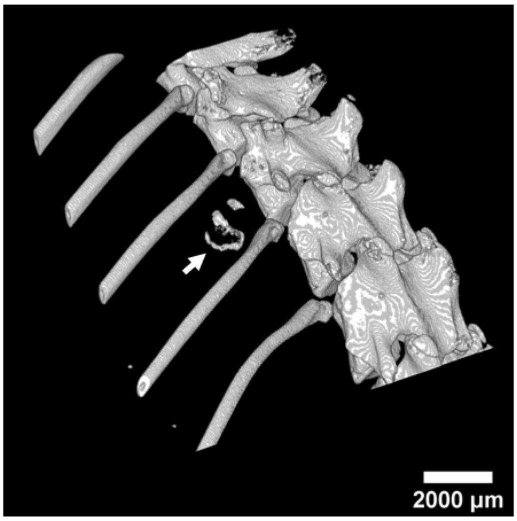


**Figure S4.** Cartilage sphere mineralization *in vivo,* 14 days after ectopic implantation in a rat model. Arrow points at implanted cartilage spheres, two spheres were implanted. Spheres were visualized with micro-computed tomography (*µ*CT).

**Supporting media:**

- **Video S1.** Three-dimensional visualization of vascular-like structures, collagen, hydroxyapatite and cell nuclei prior to implantation (D28).
- **Video S2.** Three-dimensional visualization of cell migration in defect implanted with fibrin gel (D42).
- **Video S3.** Three-dimensional visualization of cell migration in defect implanted with platelet gel (D42).
- **Video S4.** Three-dimensional visualization of cell migration in defect implanted with cartilage spheres (D42).

**Supporting references**

[1] A.L. Ponte, E. Marais, N. Gallay, A. Langonné, B. Delorme, O. Hérault, P. Charbord, J. Domenech, The In Vitro Migration Capacity of Human Bone Marrow Mesenchymal Stem Cells: Comparison of Chemokine and Growth Factor Chemotactic Activities, Stem Cells. 25 (2007) 1737–1745. https://doi.org/10.1634/stemcells.2007-0054.

[2] W. Gilbert, R. Bragg, A.M. Elmansi, M.E. McGee-Lawrence, C.M. Isales, M.W. Hamrick, W.D. Hill, S. Fulzele, Stromal cell-derived factor-1 (CXCL12) and its role in bone and muscle biology, Cytokine. 123 (2019) 154783. https://doi.org/10.1016/j.cyto.2019.154783.

[3] F. Yang, F. Xue, J. Guan, Z. Zhang, J. Yin, Q. Kang, Stromal-Cell-Derived Factor (SDF) 1-Alpha Overexpression Promotes Bone Regeneration by Osteogenesis and Angiogenesis in Osteonecrosis of the Femoral Head, Cell Physiol Biochem. 46 (2018) 2561–2575. https://doi.org/10.1159/000489684.

[4] X. Yu, Y. Huang, P. Collin-Osdoby, P. Osdoby, Stromal Cell-Derived Factor-1 (SDF-1) Recruits Osteoclast Precursors by Inducing Chemotaxis, Matrix Metalloproteinase-9 (MMP-9) Activity, and Collagen Transmigration, 2003. https://doi.org/10.1359/jbmr.2003.18.8.1404.

[5] D. qi Li, Q. long Wan, J.L. Pathak, Z. bing Li, Platelet-derived growth factor BB enhances osteoclast formation and osteoclast precursor cell chemotaxis, J Bone Miner Metab. 35 (2017) 355–365. https://doi.org/10.1007/s00774-016-0773-8.

[6] M. Zhang, W. Yu, K. Niibe, W. Zhang, H. Egusa, T. Tang, X. Jiang, The effects of platelet-derived growth factor-BB on bone marrow stromal cell-mediated vascularized bone regeneration, Stem Cells Int. 2018 (2018) 3272098.

https://doi.org/10.1155/2018/3272098.

[7] C. Maes, L. Coenegrachts, I. Stockmans, E. Daci, A. Luttun, A. Petryk, R. Gopalakrishnan, K. Moermans, N. Smets, C.M. Verfaillie, P. Carmeliet, R. Bouillon, G. Carmeliet, Placental growth factor mediates mesenchymal cell development, cartilage turnover, and bone remodeling during fracture repair, J Clin Invest. 116 (2006) 1230–1242. https://doi.org/10.1172/JCI26772.

[8] J.A. Arnott, A.G. Lambi, C.M. Mundy, H. Hendesi, R.A. Pixley, T.A. Owen, F.F. Safadi, S.N. Popoff, The Role of Connective Tissue Growth Factor (CTGF/CCN2) in Skeletogenesis, Crit Rev Eukaryot Gene Expr. 21 (2011) 43–69. https://doi.org/10.1615/critreveukargeneexpr.v21.i1.40.

[9] Q. Yang, K.P. McHugh, S. Patntirapong, X. Gu, L. Wunderlich, P. V. Hauschka, VEGF enhancement of osteoclast survival and bone resorption involves VEGF receptor-2 signaling and β3-integrin, Matrix Biol. 27 (2008) 589–599. https://doi.org/10.1016/j.matbio.2008.06.005.

[10] Y. Liu, A.D. Berendsen, S. Jia, S. Lotinun, R. Baron, N. Ferrara, B.R. Olsen, Intracellular VEGF regulates the balance between osteoblast and adipocyte differentiation, J Clin Invest. 122 (2012) 3101–3113. https://doi.org/10.1172/JCI61209.

[11] A. Grosso, M.G. Burger, A. Lunger, D.J. Schaefer, A. Banfi, N. Di Maggio, It takes two to tango: Coupling of angiogenesis and osteogenesis for bone regeneration, Front Bioeng Biotechnol. 5 (2017) 68. https://doi.org/10.3389/fbioe.2017.00068.

[12] B.O. Zhou, L. Ding, S.J. Morrison, Hematopoietic stem and progenitor cells regulate the regeneration of their niche by secreting Angiopoietin-1, Elife. 4 (2015) e05521. https://doi.org/10.7554/eLife.05521.

[13] J. Yin, G. Gong, C. Sun, Z. Yin, C. Zhu, B. Wang, Q. Hu, Y. Zhu, X. Liu, Angiopoietin 2 promotes angiogenesis in tissue-engineered bone and improves repair of bone defects by inducing autophagy, Biomed Pharmacother. 105 (2018) 932–939. https://doi.org/10.1016/j.biopha.2018.06.078.

[14] K.B.S. Paiva, J.M. Granjeiro, Matrix Metalloproteinases in Bone Resorption, Remodeling, and Repair, in: Prog Mol Biol Transl Sci, Elsevier B.V., 2017: pp. 203–303. https://doi.org/10.1016/bs.pmbts.2017.05.001.

[15] J. Wan, G. Zhang, X. Li, X. Qiu, J. Ouyang, J. Dai, S. Min, Matrix Metalloproteinase 3: A Promoting and Destabilizing Factor in the Pathogenesis of Disease and Cell Differentiation, Front Physiol. 12 (2021) 663978.

https://doi.org/10.3389/fphys.2021.663978.

[16] N. Ortega, D.J. Behonick, Z. Werb, Matrix remodeling during endochondral ossification, Trends Cell Biol. 14 (2004) 86–93. https://doi.org/10.1016/j.tcb.2003.12.003.

[17] L. Matilla, C. Roncal, J. Ibarrola, V. Arrieta, A. Garciá-Penã, A. Fernández-Celis, A. Navarro, V. Álvarez, A. Gainza, J. Orbe, V. Cachofeiro, G. Zalba, R. Sádaba, J.A. Rodríguez, N. López-Andrés, A Role for MMP-10 (Matrix Metalloproteinase-10) in Calcific Aortic Valve Stenosis, Arterioscler Thromb Vasc Biol. (2020) 1370–1382. https://doi.org/10.1161/ATVBAHA.120.314143.

[18] L. Mao, M. Yano, N. Kawao, Y. Tamura, K. Okada, H. Kaji, Role of matrix matelloproteinase-10 in the BMP-2 inducing osteoblastic differentiation, Endocr J Adv Publ. 60 (2013) 1309–1319. https://doi.org/10.1507/endocrj.EJ13-0270.

[19] B.F. Boyce, L. Xing, Functions of RANKL/RANK/OPG in bone modeling and remodeling, Arch Biochem Biophys. 473 (2008) 139–146. https://doi.org/10.1016/j.abb.2008.03.018.

[20] Y.X. Fu, J.H. Gu, Y.R. Zhang, X.S. Tong, H.Y. Zhao, Y. Yuan, X.Z. Liu, J.C. Bian, Z.P. Liu, Osteoprotegerin influences the bone resorption activity of osteoclasts, Int J Mol Med. 31 (2013) 1411–1417. https://doi.org/10.3892/ijmm.2013.1329.

[21] D. Rodriguez, T. Thula-Mata, E. Toro, Y. Yeh, C. Holt, L. Holliday, L. Gower, Multifunctional role of osteopontin in directing intrafibrillar mineralization of collagen and activation of osteoclasts, Acta Biomater. 10 (2014) 494–507. https://doi.org/10.1016/j.actbio.2013.10.010.

[22] K. Terai, T. Takano-Yamamoto, Y. Ohba, K. Hiura, M. Sugimoto, M. Sato, H. Kawahata, N. Inaguma, Y. Kitamura, S. Nomura, Role of Osteopontin in Bone Remodeling Caused by Mechanical Stress, J Bone Miner Res. 14 (1999) 839–849. https://doi.org/10.1359/jbmr.1999.14.6.839.

[23] X. Lin, S. Patil, Y.G. Gao, A. Qian, The Bone Extracellular Matrix in Bone Formation and Regeneration, Front Pharmacol. 11 (2020) 757. https://doi.org/10.3389/fphar.2020.00757.

[24] P.K. Suen, L. Qin, Sclerostin, an emerging therapeutic target for treating osteoporosis and osteoporotic fracture: A general review, J Orthop Transl. 4 (2016) 1–13. https://doi.org/10.1016/j.jot.2015.08.004.

[25] O. Duchamp de Lageneste, C. Colnot, Periostin in bone regeneration, in: A. Kudo

(Ed.), Periostin, 2019: pp. 49–59.

[26] A. Gramoun, N. Azizi, J. Sodek, J.N.M. Heersche, I. Nakchbandi, M.F. Manolson, Fibronectin inhibits osteoclastogenesis while enhancing osteoclast activity via nitric oxide and interleukin-1β-mediated signaling pathways, J Cell Biochem. 111 (2010) 1020–1034. https://doi.org/10.1002/jcb.22791.

[27] Seiji Goda, Hiroshi Hayashi, Yosuke Ujii, Osamu Takeuchi, Reiko Komasa, Eisuke Domae, Kazuyo Yamamoto, Naoyuki Matsumoto, Takashi Ikeo, Fibronectin inhibited RANKL-induced differentiation into osteoclast, J Oral Tissue Engin. 11 (2014) 227–233. https://doi.org/10.11223/jarde.11.227.

[28] A.B. Faia-Torres, T. Goren, T.O. Ihalainen, S. Guimond-Lischer, M. Charnley, M. Rottmar, K. Maniura-Weber, N.D. Spencer, R.L. Reis, M. Textor, N.M. Neves, Regulation of human mesenchymal stem cell osteogenesis by specific surface density of fibronectin: A gradient study, ACS Appl Mater Interfaces. 7 (2015) 2367–2375. https://doi.org/10.1021/am506951c.
